# Inflammation durably imprints memory CD4+ T cells

**DOI:** 10.1101/2022.11.15.516351

**Authors:** Sophie L. Gray-Gaillard, Sabrina Solis, Han M. Chen, Clarice Monteiro, Grace Ciabattoni, Marie I. Samanovic, Amber R. Cornelius, Tijaana Williams, Emilie Geesey, Miguel Rodriguez, Mila Brum Ortigoza, Ellie N. Ivanova, Sergei B. Koralov, Mark J. Mulligan, Ramin Sedaghat Herati

## Abstract

Adaptive immune responses are induced by vaccination and infection, yet little is known about how CD4+ T cell memory differs when primed in these two contexts. Notably, viral infection is generally associated with higher levels of systemic inflammation than is vaccination. To assess whether the inflammatory milieu at the time of CD4+ T cell priming has long-term effects on memory, we compared Spike-specific memory CD4+ T cells in 22 individuals around the time of the participants’ third SARS-CoV-2 mRNA vaccination, with stratification by whether the participants’ first exposure to Spike was via virus or mRNA vaccine. Multimodal single-cell profiling of Spike-specific CD4+ T cells revealed 755 differentially expressed genes that distinguished infection- and vaccine-primed memory CD4+ T cells. Spike-specific CD4+ T cells from infection-primed individuals had strong enrichment for cytotoxicity and interferon signaling genes, whereas Spike-specific CD4+ T cells from vaccine-primed individuals were enriched for proliferative pathways by gene set enrichment analysis. Moreover, Spike-specific memory CD4+ T cells established by infection had distinct epigenetic landscapes driven by enrichment of IRF-family transcription factors, relative to T cells established by mRNA vaccination. This transcriptional imprint was minimally altered following subsequent mRNA vaccination or breakthrough infection, reflecting the strong bias induced by the inflammatory environment during initial memory differentiation. Together, these data suggest that the inflammatory context during CD4+ T cell priming is durably imprinted in the memory state at transcriptional and epigenetic levels, which has implications for personalization of vaccination based on prior infection history.

**One Sentence Summary:** SARS-CoV-2 infection versus SARS-CoV-2 mRNA vaccination prime durable transcriptionally and epigenetically distinct Spike-specific CD4+ T cell memory landscapes.

## Introduction

T cell memory is crucial for durable protection against viral infection and is a well-established correlate of immune protection^1–5^. In particular, CD4+ T cells contribute to multifaceted defense via coordination of innate immune responses, help to B cells and CD8+ T cells, and direct interaction with infected cells^6, 7^. The quality of memory CD4+ T cell responses is typically assessed by interrogation of cellular frequency, cytokine production, provision of help, and T cell receptor (TCR) affinity for antigen^8–11^. The goal of vaccination is to induce long-lasting protection, however the factors that contribute to an “optimal” memory CD4+ T cell response are not fully-understood.

Following the onset of the COVID-19 pandemic, some individuals developed CD4+ T cell memory to Spike protein first through infection and others through SARS-CoV-2 vaccination. Indeed, SARS-CoV-2 infection and mRNA vaccination induced durable and robust Spike-specific CD4+ T cell responses skewed towards a Th1 and T follicular helper (T_FH_) profile^12–17^. Moreover, infection- and vaccination-derived Spike-specific CD4+ T cells rapidly expand after Spike protein exposure in subsequent SARS-CoV-2 vaccination or infection, demonstrating effective recall of memory CD4+ T cells^18–20^. In short, mRNA vaccination and SARS-CoV-2 infection each induced comparable frequencies of memory CD4+ T cell responses.

Aside from quantitative induction of a memory CD4+ T cell response by vaccine or virus, little is known about whether, or even if, infection- and vaccine-derived memory CD4+ T cells qualitatively differ, given differences in the inflammatory context in which Spike epitopes are presented. Factors during initial priming such as inflammatory signals, site and persistence of antigen exposure, cell-to-cell interactions, and cytokine milieu imprint resulting memory pools and influence T cell responses upon antigen re-exposure^21, 22^. Notably, viral infection triggers a widespread high state of inflammation in contrast to that present during immunization^23^. Indeed, COVID-19 patient serum levels of inflammatory cytokines positively correlated with disease severity when compared to healthy controls^24–26^. In contrast, serum levels of inflammatory cytokines were low following vaccination^27^ and only mild reactogenicity was reported^28–30^. The presence of concomitant inflammation may have deleterious consequences for T cell memory. Indeed, reduced TCR diversity, decreased frequency of antigen-specific memory T cells, and impaired cytokine production have been reported in the setting of excess inflammation^31–33^. How infection-associated inflammation affects Spike-specific memory CD4+ T cells has not been studied, but examination of the direct effects of inflammation on memory T cell quality will improve our understanding of infection-derived protective immunity for SARS-CoV-2 and broadly inform new strategies for optimal vaccine design.

Due to the difference in inflammatory context between infection and vaccination at the time of memory CD4+ T cell priming, we hypothesized that qualitative differences in Spike-specific memory CD4+ T cells may have been established. To test this hypothesis, we explored the transcriptional profiles of Spike-specific CD4+ T cell responses pre- and post-third vaccine dose, referenced as the “booster” dose, in individuals whose first exposure to Spike protein was either infection (infection-primed) or immunization (vaccine-primed). Using multimodal single-cell RNA sequencing (scRNAseq) following overnight Spike peptide pool stimulation in the activation-induced markers (AIM) assay^12, 34–37^, we found that Spike-specific CD4+ T cells from infection-primed individuals following the booster exhibited an increased cytotoxic signature compared to vaccine-primed individuals. Although both vaccine-primed and infection-primed participants had received the two-dose primary mRNA vaccination series, we uncovered greater expression of interferon-stimulated genes (ISGs) and interferon-related pathways in Spike-specific CD4+ T cells from infection-primed adults. In contrast, proliferative pathways were enriched in Spike-specific CD4+ T cells from vaccine-primed participants receiving the booster. Next, we compared the epigenetic landscape of Spike-specific CD4+ T cells six months post-infection and post-vaccination and found distinct chromatin patterns in each cohort driven by the IRF family of transcription factors. To test whether Spike-specific memory CD4+ T cells from vaccine-primed individuals could be affected by subsequent inflammation, we evaluated the same individuals who later had breakthrough infection. We found that breakthrough infection modestly altered the transcriptional profile of vaccine-primed participants but did not introduce the ISG-associated gene signatures seen in Spike-specific CD4+ T cells from infection-primed individuals. These data suggest a durable, inflammatory imprint on memory CD4+ T cells primed by viral infection and highlight the importance of understanding the inflammatory context of initial antigen exposure during memory CD4+ T cell development.

## Results

### Spike-specific CD4+ T cells form a distinct cluster in the activation-induced marker assay

SARS-CoV-2 infection and mRNA vaccination induced Spike-specific memory CD4+ T cell responses, as assessed by ELISpot, tetramer, and activation-induced marker (AIM) assay^13–17, 38–40^. Given that the magnitude of ISG activity differed during the acute response to vaccination and infection^23, 26, 27^, we sought to understand the extent to which concomitant inflammation affected the generation of Spike-specific memory CD4+ T cells and subsequent responses following re-exposure to Spike antigens.

Since 2020, we have been longitudinally following individuals after diagnosis of acute COVID-19 or after initiation of mRNA vaccination, as previously described^12, 41^. From this cohort, we randomly selected 22 individuals with samples available from around 8 months after second vaccination, referred to as the pre-booster time point, and one month post-booster to longitudinally assess immune memory (**Fig. 1A, S1A**). Of these, eleven did not have COVID-19, as confirmed by N antibody ELISA at the time of initial mRNA vaccination, and thus their first exposure to Spike protein was mRNA vaccination (labeled vaccine-primed). In addition, eleven individuals had confirmed diagnosis of SARS-CoV-2 infection in the spring of 2020 (labeled infection-primed), then received two doses of mRNA vaccination ∼9 months later, and received their third vaccination around 20 months after onset of COVID-19 (**Table S1-3**). Of these eleven infection-primed participants, ten had mild or minimally symptomatic COVID-19 and one had severe disease (**Table S2**). Participants’ ages ranged from 28 - 62, with a median age of 42.5 for infection-primed individuals and 39 for vaccine-primed individuals (**Table S1**).

**Figure 1.**
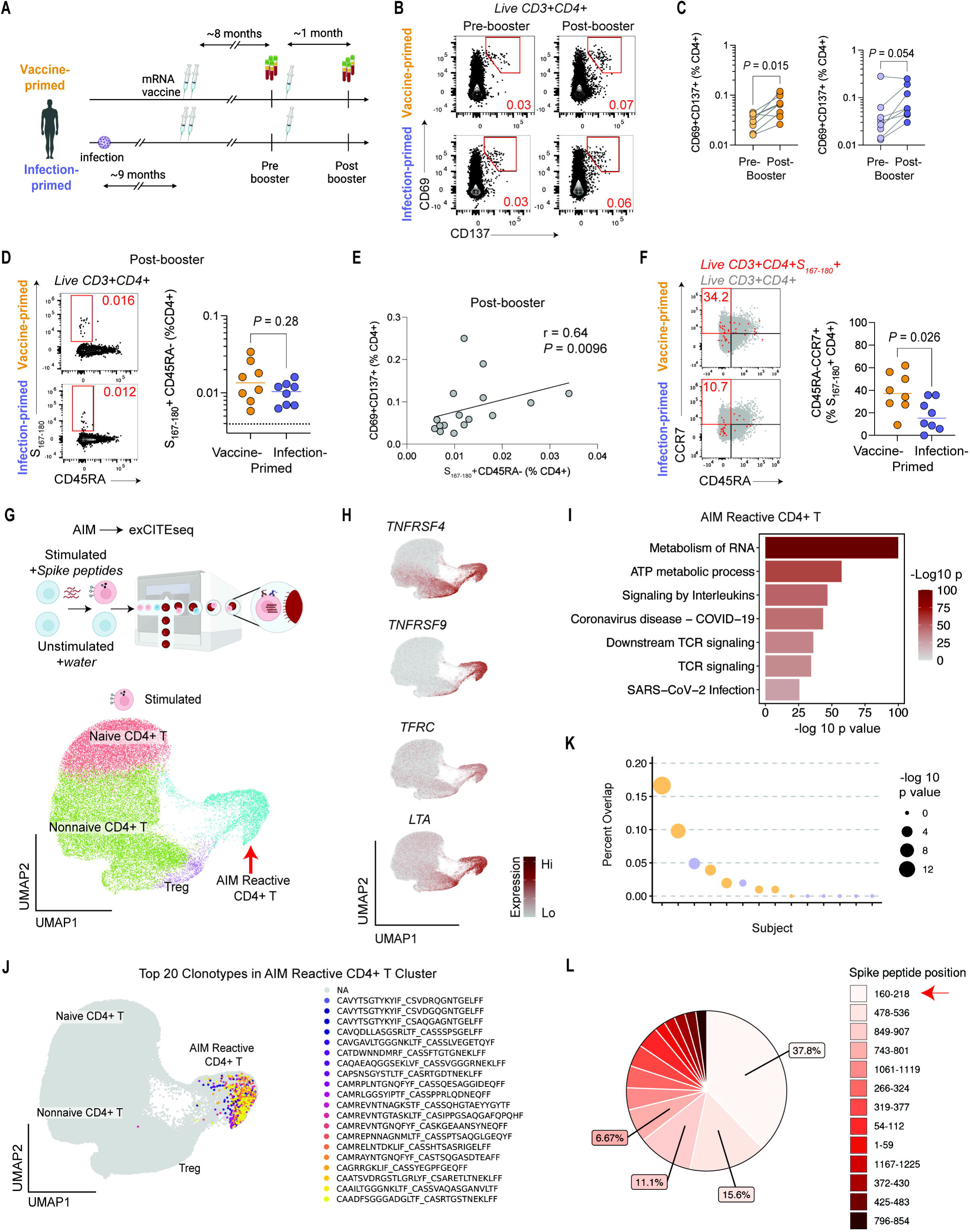
Spike-specific CD4+ T cells form a distinct cluster in the activation-induced marker assay. **A**. Study schematic. **B**. Representative flow cytometry plots for vaccine- and infection-primed participants at pre-booster and post-booster time points for expression of CD69 and CD137 after the activation-induced markers (AIM) assay. Frequency shown in red. **C**. Summary data for vaccine-primed (orange, n = 8) and infection-primed (purple, n=8) participants’ PBMC after AIM assay. *P* values by Wilcoxon matched-pairs signed rank test. **D**. S_167-180_ HLA class II tetramer and CD45RA expression plots for vaccine- and infection-primed participants at the post-booster time point. Summary data shown for vaccine-primed (orange, n = 8) and infection-primed (purple, n=8) participants. *P* value by Wilcoxon test. Frequency shown in red. **E**. Correlation between frequency of CD69+CD137+ CD4+ T cells after AIM and the frequency of S_167-180_+ CD4+ T cells at the post-booster time point (n = 16). **F**. Representative plots for CD45RA and CCR7 expression by S_167-180_+ CD4+ T cells from vaccine- and infection-primed participants. *P* value by Wilcoxon test. Frequency shown in red. **G**. Schematic of experimental design. UMAP projection of CD4+ T cells from the AIM assay, pooled across samples and clustered using gene expression. **H**. Scaled expression of *TNFRSF4* (OX40), *TNFRSF9* (CD137), *TFRC* (CD71), *LTA*. **I**. Gene ontology for differentially genes expressed at adjusted *P* < 0.05 for AIM-Reactive CD4+ T cell compared to CD4+ T cell cluster. **J**. Top 20 clonotypes identified among the AIM-Reactive CD4+ T cluster. **K**. Per-participant TCR overlap between AIM-Reactive CD4+ TCRs and the Adaptive Biotechnologies SARS-CoV-2-reactive TCR database. **L**. Distribution of epitope specificity of AIM-Reactive CD4+ T cells as inferred from the reference SARS-CoV-2 TCRβ dataset. See also Figure S1 and Tables S1-S2.

To identify vaccine- and infection-primed participants’ Spike-specific memory CD4+ T cells, we used the AIM assay^12, 34–37, 42^ in which PBMCs are stimulated overnight with overlapping Spike peptide pools and evaluated for surface protein expression. AIM-reactive CD4+ T cells were identified based on co-expression of activation markers such as CD69, CD137, and CD200 (**Fig. 1B, S1B)**. Stimulated cells demonstrated higher frequencies of CD69+ CD137+ CD4+ T cells than paired unstimulated controls (*P* < 0.0001; n=18; Wilcoxon matched-pairs signed rank test) (**S1C**). Both cohorts demonstrated an increase in Spike-specific CD4+ T cells after booster (**Fig 1C, S1D**), suggesting that a third dose of mRNA vaccination indeed elicited a CD4+ T cell response, as reported previously^43, 44^.

We next probed epitope-specific CD4+ T cell responses using an HLA-DPB1*04:01 allele tetramer loaded with peptide 167-180 of Spike protein (TFEYVSQPFLMDLE, S_167–180_) as reported previously^38^. We interrogated circulating S_167-180_-specific CD45RA- CD4+ T cells one month after booster vaccination and observed similar frequencies between cohorts (n=8 each for vaccine- and infection-primed cohorts; *P*-value = 0.28, Wilcoxon test) (**Fig. 1D, S1E**). Moreover, the frequency of S_167-180_-specific CD45RA-CD4+ T cells positively correlated with AIM responding CD4+ T cells measured by flow cytometry (**Fig 1E, S1F**). These results demonstrated that the AIM-reactive CD4+ T cells were strongly enriched for Spike-specific memory CD4+ T cells in all vaccine- and infection-primed participants.

We observed subtle differences between vaccine- and infection-primed Spike-specific memory CD4+ T cells by surface protein expression (**Fig. 1E**). To more deeply profile the differences between the Spike-specific memory CD4+ T cells in vaccine- and infection-primed participants, we evaluated the transcriptional state of Spike-specific CD4+ T cells using droplet-based multimodal scRNAseq^45^ for 14 randomly selected participants (**Fig. S1A**) following AIM assay and subsequent magnetic bead enrichment for CD69 and CD137, two surface markers highly expressed by AIM-reactive cells^12, 37^ (**Fig. 1B**). High-quality single cells from 33 samples were integrated for downstream analyses (**Fig. S1G**). In total, we recovered 127,047 CD4+ T cells. Dimension reduction was performed on gene expression to generate a uniform manifold approximation and projection (UMAP). Using graph-based clustering^46^, we identified 4 major clusters, one of which was nearly absent in unstimulated controls (**Fig. 1G, S1H**). We denoted this cluster of 11,919 cells as AIM-Reactive CD4+ T. We then interrogated genes associated with T cell activation^36^. We found upregulation of genes such as *TNFRSF9* (CD137), consistent with the magnetic bead enrichment performed. Genes such as *TNFRSF4* (OX40) and *TFRC* (CD71) were also upregulated in the AIM-Reactive CD4+ T cluster (**Fig. 1H**). Additionally, the AIM-Reactive CD4+ T cluster enriched for *IFNG*, *IL2*, and *LTA* transcripts (**Fig. S1I**). Gene ontology analysis of genes differentially expressed by AIM-Reactive CD4+ T cluster compared to the Nonnaive CD4+ T cluster demonstrated enrichment for terms such as “cellular response to stimuli” and “TCR signaling” (**Fig. 1I**). Together, these data demonstrated robust identification of the AIM-reactive subset of CD4+ T cells by scRNAseq.

To further test whether AIM-reactive CD4+ T cells were Spike-specific, we investigated TCR sequences. The clonal AIM-reactive TCRs were not observed in any other cluster (**Fig. 1J**). Next, we compared the AIM-Reactive CD4+ T cell cluster TCRβ sequences to an independently-generated database of ∼3000 Class II TCRβ^47^. For eight of 14 participants, there was substantial overlap between the AIM-reactive CD4+ T cell CDR3 amino acid sequences and those of the public SARS-CoV-2 TCRβ database^47^ (**Fig. 1K**). TCR overlap was not observed in all participants, perhaps due to the relatively limited number of TCRβ sequences available for comparison. The TCRβ sequences that were congruent between our dataset and the public database were evaluated for epitope specificity^47^. As was previously seen with Spike-specific CD4+ T cells found in human lymph nodes after SARS-CoV-2 vaccination^38^, the majority of TCRs in the AIM-Reactive cluster mapped to peptides 160-218 of Spike protein (**Fig. IL**), which was reported to contain immunodominant CD4+ T cell epitopes^39^. In sum, multimodal single cell profiling of AIM-Reactive cells could be used to more deeply evaluate Spike-specific memory CD4+ T cells.

### Infection-primed Spike-specific CD4+ T cells exhibit a cytotoxic profile

Prior studies in the literature described expression of cytokines including IFNg, TNF, IL-2, and granzyme B following SARS-CoV-2 infection and vaccination^12–14, 17, 18, 39, 48^. Moreover, cytotoxic Spike-specific memory CD4+ T cells are formed following SARS-CoV-2 infection^49–51^. We found that overnight stimulation with Spike peptides induced more transcripts for these cytokines in the AIM-Reactive CD4+ T cluster relative to non-activated CD4+ T cells (**Fig. 2A, Fig. S2A**). To investigate whether vaccine- and infection-primed Spike-specific memory CD4+ T cells demonstrated different functionality, we compared the expression of *IFNG, TNF, IL2,* and *GZMB* transcripts. Of these, *IL2* expression was higher for vaccine-primed participants, whereas *GZMB* expression was higher in the infection-primed cohort, at both the pre- and post-booster time points (**Fig. 2B**). As polyfunctional CD4+ T cell responses are a correlate of protection^10^ and have been evaluated following acute COVID-19 or SARS-CoV-2 vaccination^27, 48, 52^, we assessed differences in polyfunctionality in AIM-reactive CD4+ T cells from vaccine- and infection-primed cohorts at both time points. Spike-specific CD4+ T cell polyfunctionality was largely similar across cohorts, with little change due to booster immunization (**Fig. 2C, S2B**). Infection-primed individuals had higher frequency of *IFNG+TNF+GZMB+* cells than their vaccine-primed counterparts at both pre-booster (*P* = 0.047, Wilcoxon test) and post-booster (*P* = 0.006, Wilcoxon test) time points. In contrast, vaccine-primed participants had a higher frequency of *TNF*+*IL2*+ Spike-specific CD4+ T cells post-booster (*P* = 0.004, Wilcoxon test) (**Fig. S2B**).

**Figure 2.**
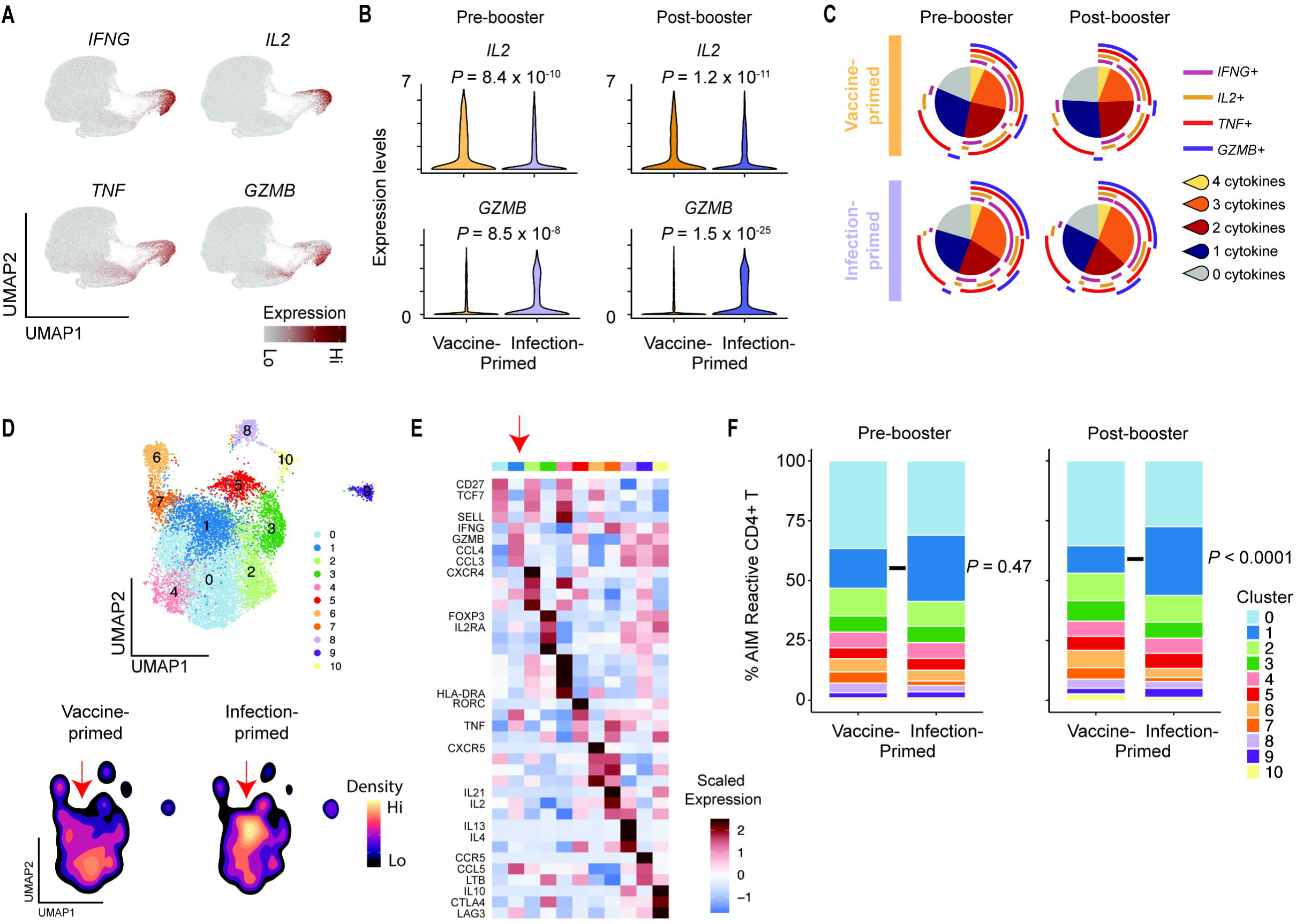
Infection-primed Spike-specific CD4+ T cells exhibit a cytotoxic profile. **A**. Scaled expression of *IFNG*, *IL2*, *TNF,* and *GZMB*. **B**. Scaled expression for *IL2* and *GZMB* between cohorts at pre- and post-booster timepoints. Nominal *P* values reported. **C**. Polyfunctionality analysis. **D**. Spike-specific CD4+ T cells projected onto UMAP and clustered for gene expression of 27 select parameters. UMAP colored by density and split by cohort. **E**. Scaled expression of differentially expressed genes at adjusted *P* < 0.01 for each cluster. **F**. Vaccine-primed versus infection-primed Spike-specific CD4+ T cell cluster distribution at pre- and post-booster time points (unpaired two-way ANOVA with Sidak posttest). See also Figure S2.

Others have reported Th1 and T_FH_ subsets following SARS-CoV-2 infection and vaccination^17, 18, 38^. To evaluate if Spike-specific CD4+ T cells differed in subset distribution, we applied dimensionality reduction with 27 parameters to the AIM-Reactive CD4+ T cluster which revealed 11 unique clusters (**Fig. 2D**). The distribution of cells across the 11 clusters were similar between pre- and post-booster timepoints for both cohorts (**Fig. S2C**). Cluster 0, which was enriched for strongly expressed *TCF7*, *CD27*, and *SELL*, contained the majority of Spike-specific CD4+ T cells and was similar between cohorts (**Fig 2E**). However, cluster 1, which enriched for *GZMB*, *CCL3, CCL4* as well as *PRF1*, *GZMH*, *NKG7*, was comprised of more Spike-specific CD4+ T cells from infection-primed than from vaccine-primed participants following booster vaccination (*P* < 0.0001, two-way ANOVA, **Fig 2D-F, S2D**). These data demonstrated that infection-primed Spike-specific CD4+ T cells had an augmented cytotoxic profile compared to vaccine-primed Spike-specific CD4+ T cells. Our findings indicate maintenance of functional capability after repeated mRNA vaccinations, including in infection-primed participants.

### Inflammation during priming results in durable transcriptional effects in Spike-specific CD4+ T cells

Given the differences observed between the cohorts using a subset-focused analysis (**Fig 2**), we next asked whether the broader transcriptional profile of Spike-specific CD4+ T cells differed by mechanism of priming. Principal component analysis (PCA) of the post-booster samples revealed separation between vaccine- and infection-primed participants (**Fig 3A**). To identify the genes driving the differences between cohorts, we performed differential gene expression analysis of the AIM-Reactive CD4+ T cluster before and after booster immunization. Between Spike-specific memory CD4+ T cells from vaccine- and infection-primed participants, there were 317 genes differentially expressed pre-booster and 755 genes differentially expressed post-booster (**Fig. S3A-B, Table S4**), which were not observed in analysis of other clusters (**Fig S3C**). Spike-specific CD4+ T cells from vaccine-primed individuals differentially expressed genes such as *NFKBID, NFKBIA, NFKB2*, *NFKBIZ*, and *REL,* which may reflect differential NF-kB signaling. Additionally, *CCR7* expression was enriched in vaccine-primed Spike-specific CD4+ T cells at both time points, which corroborated our previous observation of differential surface protein expression of CCR7 by flow cytometry in S_167-180_+ CD4+ T cells (**Fig. 1E**). However, *CCR7* expression, as a proxy for differentiation state of the memory CD4+ T cells, did not drive the observed differences between the cohorts (**Fig S3D**). In contrast, Spike-specific CD4+ T cells from infection-primed individuals differentially expressed genes such as *HLA-B*, *GZMB*, *IFITM1*, *IFITM3*, and *IFI6*, which are ISGs^53^ (**Fig. 3B**). Thus, these data revealed distinct transcriptional profiles between vaccine- and infection-primed Spike-specific CD4+ T cells.

**Figure 3.**
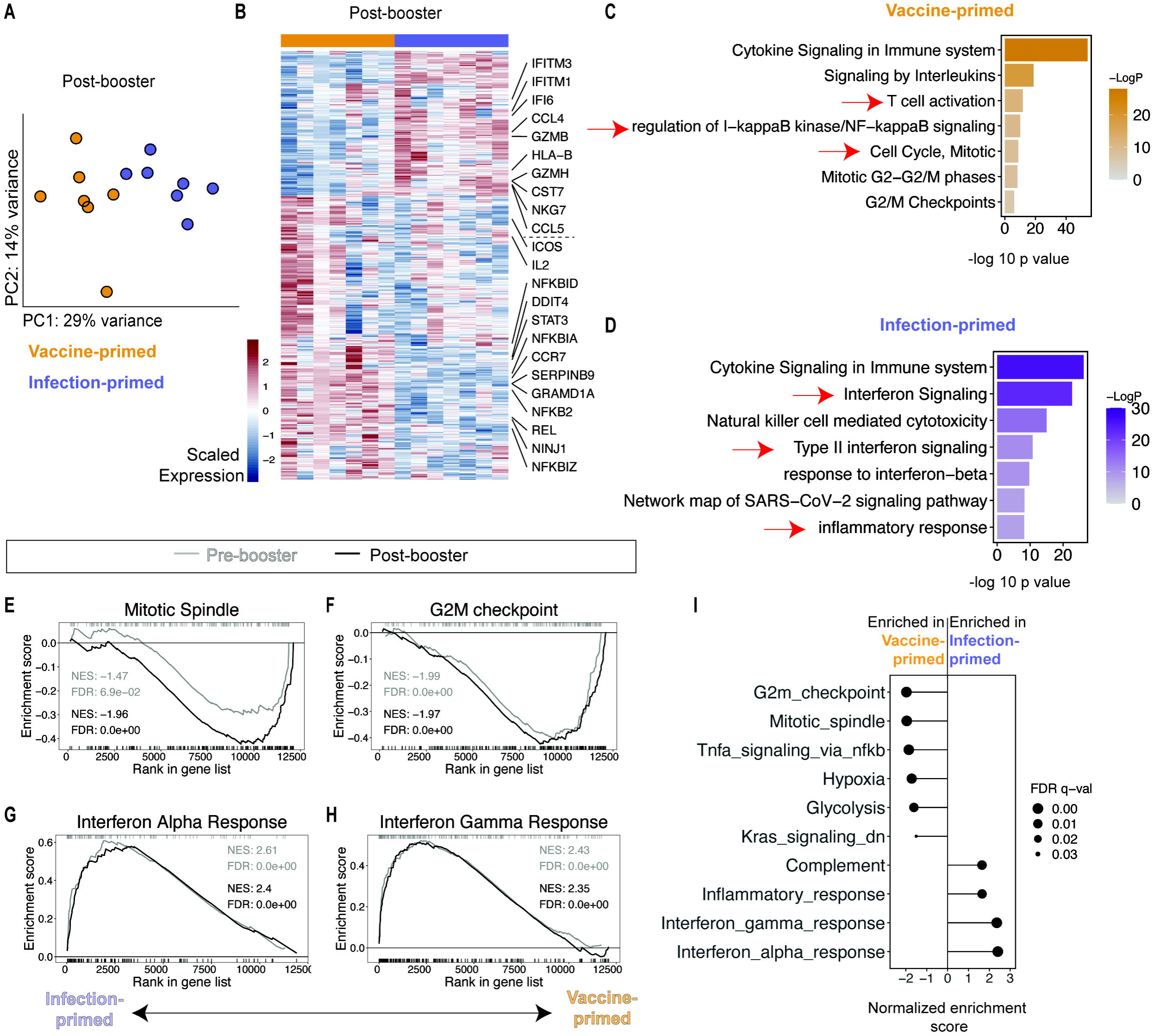
Inflammation during priming results in durable transcriptional effects in Spike- specific CD4+ T cells. **A**. PCA of post-booster vaccine-primed (orange) and infection-primed (purple) Spike-specific CD4+ T cells. Each symbol indicates one participant. **B**. Heatmap of differentially expressed genes between post-booster vaccine- and infection-primed cohorts. **C-D**. Gene ontology analysis for differentially genes expressed at adjusted *P* < 0.05 for vaccine-primed (**C**) and infection-primed (**D**) Spike-specific CD4+ T cells at post-booster time point. **E-H**. GSEA for Mitotic Spindle (**E**), G2M Checkpoint (**F**), Interferon Alpha (**G**), and Interferon Gamma Response (**H**) gene sets for Spike-specific CD4+ T cells. Light gray line indicates pre-booster time point and dark gray line indicates post-booster time point. **I**. GSEA results for Hallmark gene sets enriched at *FDR* < 0.05 post-booster. Positive enrichment scores denote enrichment towards the infection-primed cohort in (**E-I**). See also Figure S3 and Tables S4-S5.

We next evaluated the coordinated gene expression of the vaccine-primed Spike-specific CD4+ T cells. Gene ontology analysis of upregulated genes in Spike-specific CD4+ T cells from the vaccine-primed cohort at the post-booster time point (**Fig 3B, S3B**) revealed terms for “Cell Cycle” and “NF-kappa B signaling” (**Fig 3C-D**). To test whether there were differences between cohorts at a pathway level, we evaluated the coordinated expression of genes using gene set enrichment analysis (GSEA)^54^. There was strong enrichment for Mitotic Spindle and G2M Checkpoint signatures in the vaccine-primed Spike-specific CD4+ T cells at the pre- and post-booster time points (**Fig 3E-F, S3C**), as well as enrichment for the NF-kB signaling gene set in the Spike-specific CD4+ T cells from vaccine-primed participants at the post-booster time point (**Fig 3I, S3C, Table S5**). These data suggested greater expression of genes related to proliferation in the vaccine-primed Spike-specific CD4+ T cells and may have implications for subsequent cellular activation and proliferative potential.

In the Spike-specific CD4+ T cells from the infection-primed cohort, gene ontology analysis of differentially expressed genes (DEGs) (**Fig 3B, S3B**) denoted strong enrichment of terms for “interferon signaling” and “inflammatory response” (**Fig. 3D**). Additionally, GSEA revealed enrichment for Interferon Alpha Response and Interferon Gamma Response gene sets in Spike-specific CD4+ T cells from infection-primed individuals at the pre- and post-booster time points (**Fig 3G-H, S3C, Table S5**), consistent with the gene ontology analysis (**Fig. 3D**). Several other gene sets were differentially enriched, including Inflammatory Response and Complement in infection-primed individuals (**Fig 3I, S3C**). Furthermore, booster immunization did not substantially change gene set enrichment (**Fig. 3E-H, S3C**). All together, these findings suggested that Spike-specific CD4+ T cells were differentially imprinted at the time of priming and that the transcriptional profile during reactivation changed minimally following subsequent Spike protein exposure by mRNA vaccination.

### Vaccine-primed and infection-primed Spike-specific CD4+ T cells have distinct epigenetic landscapes

Given durability of the inflammatory imprint across time and Spike re-exposure (**Fig 3**), we next wanted to evaluate whether the infection-versus vaccine-derived transcriptional signatures were encoded at an epigenetic level. We hypothesized that the inflammatory conditions during initial priming would result in distinct patterns of chromatin accessibility in Spike-specific memory CD4+ T cells. Thus, PBMCs were analyzed from individuals at least 6 months after priming by either acute COVID-19 (n=6) or by primary mRNA vaccine series (n=6) (**Fig. 4A, S4A, Table S2**).

**Figure 4.**
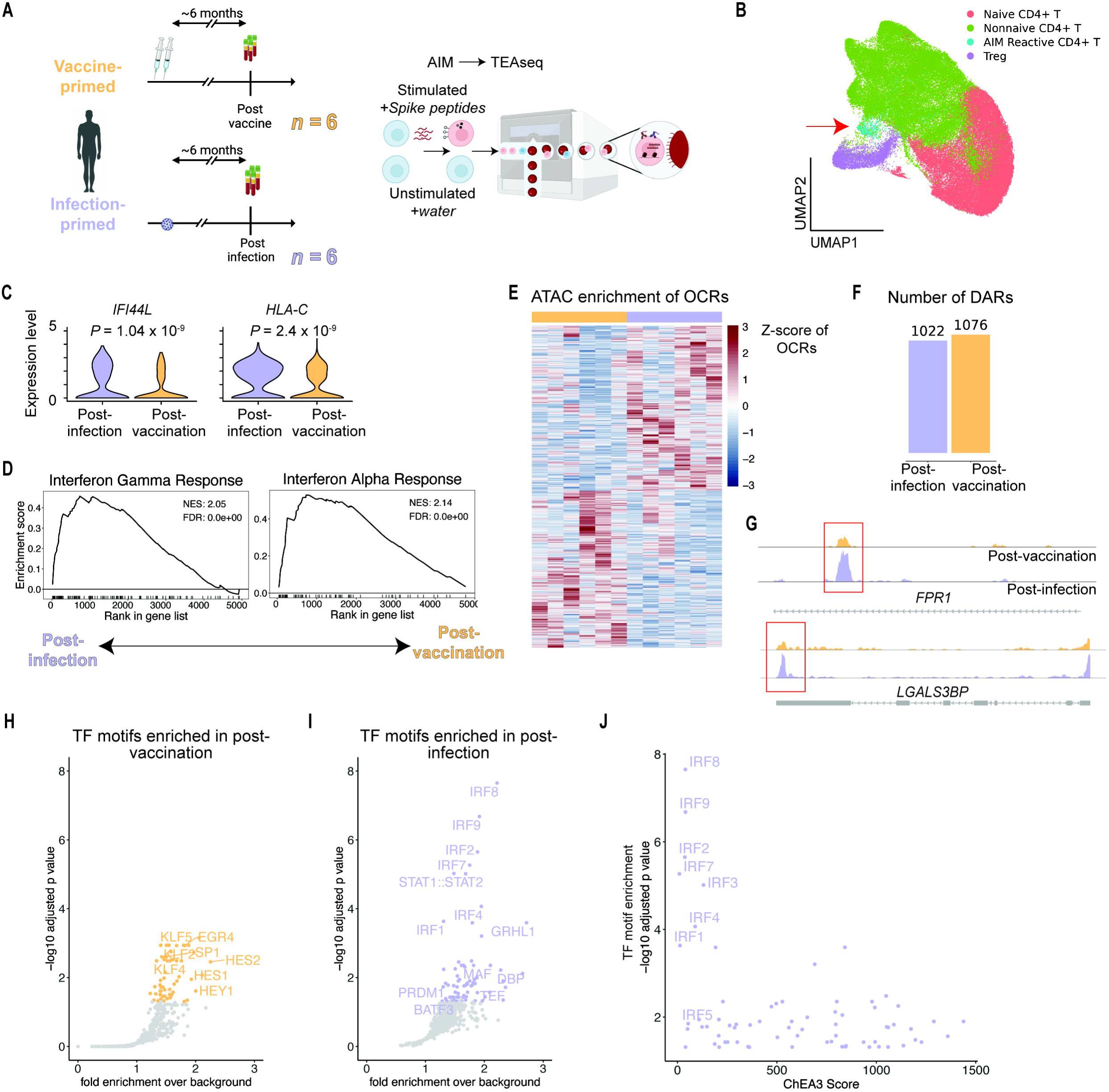
Vaccine-primed and infection-primed Spike-specific CD4+ T cells have distinct epigenetic landscapes. **A**. Study schematic. **B**. UMAP of stimulated CD4+ T cells pooled across samples and clustered for gene expression. **C**. Difference in scaled expression for *IFI44L* and *HLA-C* between post-infection and post-vaccination samples. Nominal *P* values shown and determined by DESeq2. **D**. GSEA for Interferon Gamma Response and Interferon Alpha Response gene sets for transcriptional profiling of Spike-specific memory CD4+ T cells. Positive enrichment scores indicate enrichment for the post-infection cohort. **E**. Heatmap open chromatin regions (OCRs) in the Spike-specific CD4+ T cells post-vaccination (yellow) and post-infection (purple). **F**. Number of differentially accessible regions (DARs) shown in (**E**) at a nominal *P* value < 0.05. **G**. Representative ATAC-seq tracts shown at the *FRP1* and *LGALS3BP* loci. **H-I**. Enrichment of transcription factor (TF) binding motifs over background in the top differentially-accessible OCRs based on nominal *P* value < 5×10^-3^ in post-vaccination Spike-specific memory CD4+ T cells (**H**) and post-infection Spike-specific memory CD4+ T cells (**I**). **J**. TFs shown for the −log_10_ adjusted *P* value from post-infection OCR motif analysis against the TF predicted by ChEA3 using the infection-primed gene expression data. See also Figure S4 and Tables S6-S7.

To test this hypothesis, we profiled Spike-specific CD4+ T cells via enrichment for CD4+ T cells expressing CD69 or CD137 after overnight Spike peptide stimulation using TEA-seq^55^. Similarly to exCITE-seq profiling of cells post-AIM assay, dimensionality reduction and clustering based on gene expression revealed a cluster of AIM-Reactive CD4+ T cells that differentially expressed transcripts associated with activation (**Fig. 4B, S4B-C**). Spike-specific CD4+ T cells post-infection differentially expressed ISGs such as *IFI44L* and *HLA-C* and demonstrated strong enrichment for Interferon Alpha Response and Interferon Gamma Response gene sets by GSEA, compared to Spike-specific CD4+ T cells from post-vaccination individuals (**Fig 4C-D, Table S6**). These transcriptional differences are concordant with the transcriptional profiling during boosting (**Fig 3**) and suggest imprinting during differentiation is already evident six months after priming.

Next, we interrogated whether the transcriptional imprint was due to alteration in the chromatin accessibility landscape of Spike-specific memory CD4+ T cells. After genome-wide chromatin mapping, we observed the majority of open chromatin regions (OCRs) in intergenic regions and introns (**Fig. S4D, Table S7**). To determine whether Spike-specific CD4+ T cell chromatin accessibility differed between cohorts, we performed analysis of differentially accessible regions (DAR). In total, 2,098 DARs were identified between post-infection and post-vaccination (**Fig. 4E-F**). Consistent with the transcriptional data, ISG loci had more OCRs in post-infection than in post-vaccination Spike-specific CD4+ T cells, such as was observed in the *FPR1* and *LGALS3BP* loci (**Fig 4G**). Together, these data suggested an epigenetic basis for the differential transcriptional profiles seen in Spike-specific memory CD4+ T cells following infection or vaccination.

Inflammation is a complex phenomenon that can be mediated by many proteins. To determine which signaling pathways could have been most relevant to the transcriptional profiles observed, transcription factor (TF) motif analysis of OCRs was performed. Indeed, the post-vaccination Spike-specific CD4+ T cells enriched for HES/HEY-family TFs (**Fig. 4H**), underscoring the role of Notch signaling for CD4+ T cell memory formation^56–59^. In contrast, post-infection Spike-specific CD4+ T cells had motif enrichment for interferon regulatory factor (IRF)-family TFs and STAT1::STAT2 motifs (**Fig. 4I**). Indeed, the IRF family TFs directly influence the differentiation of CD4 T cells^60–63^. Moreover, enriched motifs from OCRs were concordant with the ChEA3-predicted TF regulation^64^ derived from genes upregulated in infection-primed samples post-booster (**Fig. 4J**) and post-vaccination samples (**Fig. S4E**). Together, these data suggest that the inflammatory context during memory CD4+ T cell formation leads to epigenetic changes that could bias subsequent activation to cognate antigen.

### Breakthrough infection minimally alters the transcriptional profile of Spike-specific CD4+ T cells

The inflammation-associated transcriptional imprint in infection-derived Spike-specific memory CD4+ T cells remained largely unchanged despite multiple mRNA vaccinations (**Fig. 3**). This provoked the question of whether vaccine-derived memory CD4+ T cells would be re-imprinted with the transcriptional changes associated with inflammation after exposure to Spike protein in the context of breakthrough infection. To assess this, we examined Spike-specific CD4+ T cells one month after breakthrough infection, defined as detection of SARS-CoV-2 virus or antigen in an individual who had been vaccinated but not previously infected. Breakthrough SARS-CoV-2 infection occurred in six of the eleven vaccine-primed participants at a median of 5 months after third vaccination, during a time period when Omicron strain was predominant. For these individuals, peripheral blood was collected one month after onset of symptoms (**Fig. 5A, S1A, Table S1**). All breakthrough infections were mild and did not require hospitalization (**Table S3**).

**Figure 5.**
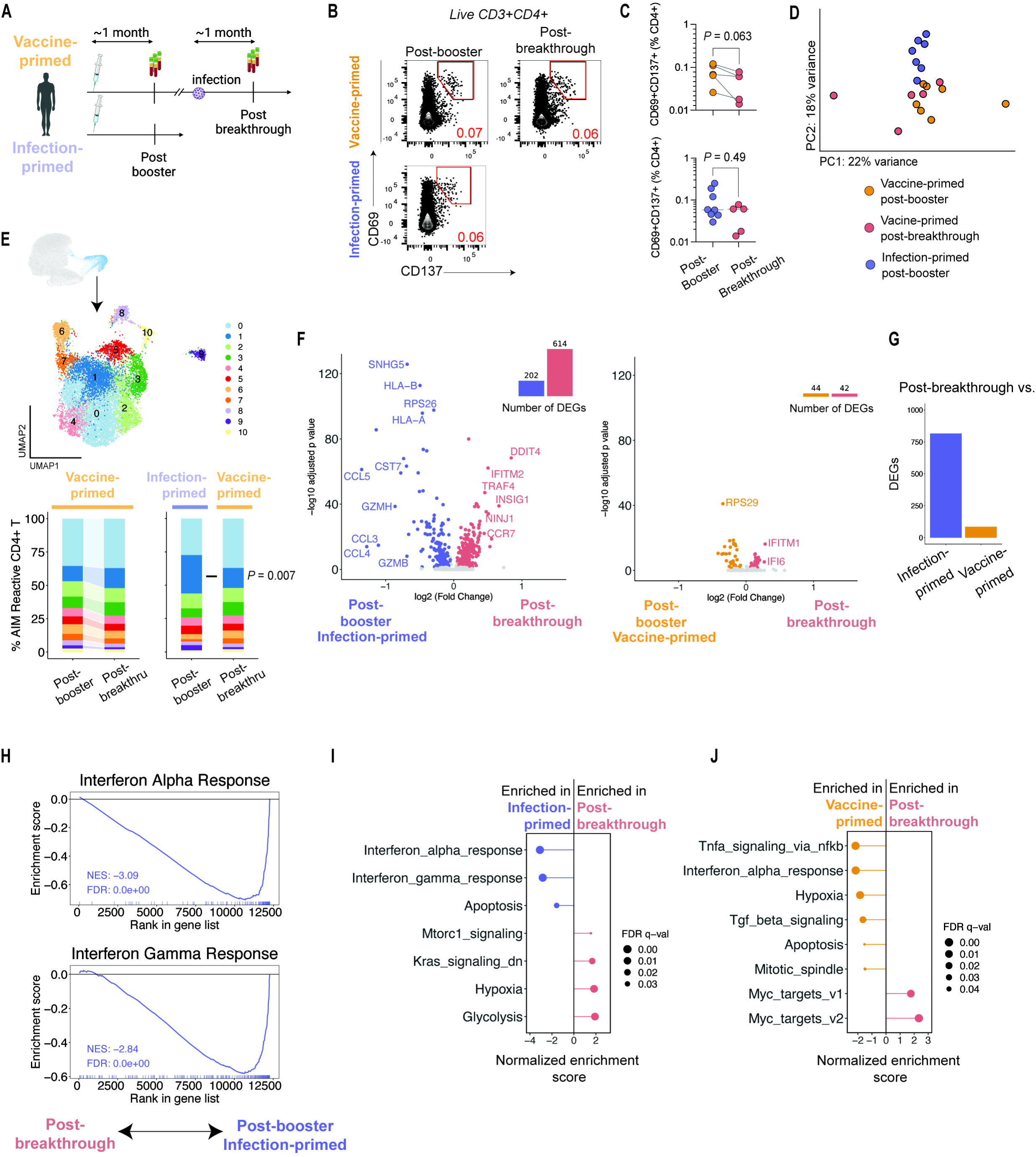
Breakthrough infection minimally alters the transcriptional profile of Spike-specific CD4+ T cells. **A**. Study schematic. **B**. Example flow plots for post-booster vaccine- and infection-primed participants and post-breakthrough samples for expression of CD69 and CD137 after AIM assay. Frequency shown in red. **C**. Summary plots of CD69+CD137+ co-expression in CD4+ T cells at post-booster and post-breakthrough time points. *P* values by Wilcoxon matched-pairs signed rank test (top) and Wilcoxon unpaired test (bottom). **D**. PCA of transcriptional profiling of post-booster vaccine-primed (orange) and infection-primed (purple) and post-breakthrough (red) Spike-specific CD4+ T cells. Each symbol indicates one participant. **E**. Spike-specific CD4+ T cells projected onto UMAP and clustered as described in **Fig 2D**. Bar graphs of vaccine-versus infection-primed Spike-specific CD4+ T cell cluster distribution at post-booster and post-breakthrough time points (two-way ANOVA). **F**. Volcano plot showing differentially expressed genes at adjusted *P* < 0.05. Genes in purple denote enrichment in infection-primed Spike-specific CD4+ T cells, orange for vaccine-primed post-booster Spike-specific CD4+ T cells, and red for post-breakthrough Spike-specific CD4+ T cells. **G**. Summary bar graph of DEGs in (**F**). **H**. GSEA for Interferon Gamma Response and Interferon Alpha Responses gene sets in infection-primed samples. **I-J.** GSEA for Hallmark gene sets at *FDR* < 0.05 when comparing infection-primed, post-booster samples to post-breakthrough (**I**) and vaccine-primed, post-booster samples to post-breakthrough (**J**). Positive enrichment scores denote enrichment for the post-breakthrough samples and negative enrichment scores signify enrichment for post-booster samples. See also Figure S5 and Tables S3, S8-S9.

We first considered Spike-specific CD4+ T cell frequencies before and after breakthrough SARS-CoV-2 infection. Indeed, prior studies described induction of Spike-specific memory CD4+ T cells in circulation, which peaked around 5-7 days after onset of symptoms and returned to pre-breakthrough frequencies by day 30^19, 20^. We assessed circulating frequency of Spike-specific CD4+ T cells using AIM one month after breakthrough SARS-CoV-2 infection and found that the post-breakthrough frequency of AIM-responding cells was comparable to matched post-booster (pre-breakthrough) samples from the same individuals (**Fig. 5B**). Moreover, frequencies of Spike-reactive CD4+ T cells following breakthrough SARS-CoV-2 infection did not differ from that of infection-primed participants (**Fig 5C**).

To test whether vaccine-primed Spike-specific memory CD4+ T cells can be transcriptionally imprinted due to concomitant inflammation, we compared transcriptional profiles for the post-booster vaccine-primed individuals, post-breakthrough vaccine-primed individuals, and post-booster infection-primed individuals, all at one month after each cohort’s re-exposure to Spike protein. Similar to what was previously observed (**Fig. 3A**), PCA once again demonstrated separation between vaccine- and infection-primed cells driven by PC2 (**Fig 5D**), indicating relative preservation of the transcriptional differences between vaccine-primed and infection-primed memory states despite breakthrough infection. Moreover, post-breakthrough vaccine-primed responses did not cluster independently of post-booster vaccine-primed samples (**Fig 5D**), suggesting transcriptional similarity between the post-booster and post-breakthrough Spike-specific CD4+ T cells from vaccine-primed individuals. We next assessed whether there was an enrichment in cytotoxic gene profiles following breakthrough infection as was seen with the infection-primed setting (**Fig. 2D**). Indeed, cluster 1 remained enriched in infection-primed Spike-specific CD4+ T cells relative to post-breakthrough vaccine-primed Spike-specific CD4+ T cells (*P* = 0.007, two-way ANOVA with Sidak post-test, **Fig. 2E**). Thus, breakthrough SARS-CoV-2 infection did not grossly alter the transcriptional profile of Spike-specific memory CD4+ T cells.

Next, we interrogated the broader transcriptional landscape of Spike-specific memory CD4+ T cells following breakthrough SARS-CoV-2 infection. A similar pattern of differential gene expression was observed when comparing post-breakthrough vaccine-primed and infection-primed Spike-specific CD4+ T cells (**Fig. 5F, Table S7**) to what was seen previously when comparing post-booster vaccine-primed and infection-primed Spike-specific CD4+ T cells (**Fig. 3**), including differential expression of *HLA-B*, *CCL5*, and *GZMB* by the infection-primed Spike-specific CD4+ T cells relative to post-breakthrough Spike-specific CD4+ T cells (**Fig. 5F)**. We next asked if post-breakthrough vaccine-primed Spike-specific CD4+ T cells were transcriptionally distinct from post-booster vaccine-primed Spike-specific CD4+ T cells, however, relatively few genes were differentially expressed between these two conditions (**Fig 5F-G**). In summary, these results suggested that breakthrough infection minimally altered the transcriptional profile of established Spike-specific memory CD4+ T cells.

We then asked if the pathway-level transcriptional changes were similar to the differential expression analysis described above. Indeed, GSEA demonstrated enrichment of Interferon Alpha Response and Interferon Gamma Response gene sets in the infection-primed Spike-specific CD4+ T cells relative to the post-breakthrough vaccine-primed Spike-specific CD4+ T cells (**Fig 5H-I, S5C, Table S9**). Moreover, GSEA did not demonstrate increased enrichment for Interferon Alpha Response in Spike-specific CD4+ T cells from post-breakthrough vaccine-primed cells, relative to post-booster vaccine-primed cells (**Fig 5J**). Together, these data demonstrate that breakthrough SARS-CoV-2 infection in individuals resulted in few alterations to the transcriptional profile of vaccine-primed Spike-specific CD4+ T cells, further supporting the notion of relative durability of the transcriptional imprinting in Spike-specific memory CD4+ T cells.

### Clonotypes enriched for the vaccine-primed gene signature can undergo inflammatory imprinting by breakthrough infection

Given the robust and durable differences observed between vaccine- and infection-primed Spike-specific memory CD4+ T cells, we wanted to examine heterogeneity in the transcriptional profiles at single cell resolution. To do this, we derived a ‘vaccine signature’ and an ‘infection signature’ from the post-booster DEGs (**Fig. S3B**) and determined each cell’s molecular imprint scores by Gene Set Variation Analysis (GSVA)^65^. Enrichment scores for the vaccine and infection imprints were plotted in 2-D Cartesian space to visualize the single cell composition of each cohort’s Spike-specific memory CD4+ T cell pool (**Fig. 6A, S6A**). The highest frequency of vaccine-primed Spike-specific CD4+ T cells was in the quadrant most positively enriched for the vaccine signature and most negatively enriched for the infection signature, whereas the highest frequency of infection-primed cells was in the quadrant most positively enriched for the infection signature and most negatively enriched for the vaccine signature (**Fig. 6B-C, S6B**). We next asked if breakthrough SARS-CoV-2 infection affected the distribution observed for vaccine-primed Spike-specific CD4+ T cells. Despite breakthrough infection, Spike-specific memory CD4+ T cells remained predominantly in the quadrant that was positively enriched for the vaccine signature and negatively enriched for the infection signature. Indeed, there were more infection-primed Spike-specific CD4+ T cells that were positively enriched for the infection signature than post-breakthrough Spike-specific CD4+ T cells (**Fig. 6D, S6C**). These data suggest the transcriptional imprinting is present across the majority of Spike-specific CD4+ T cells.

**Figure 6.**
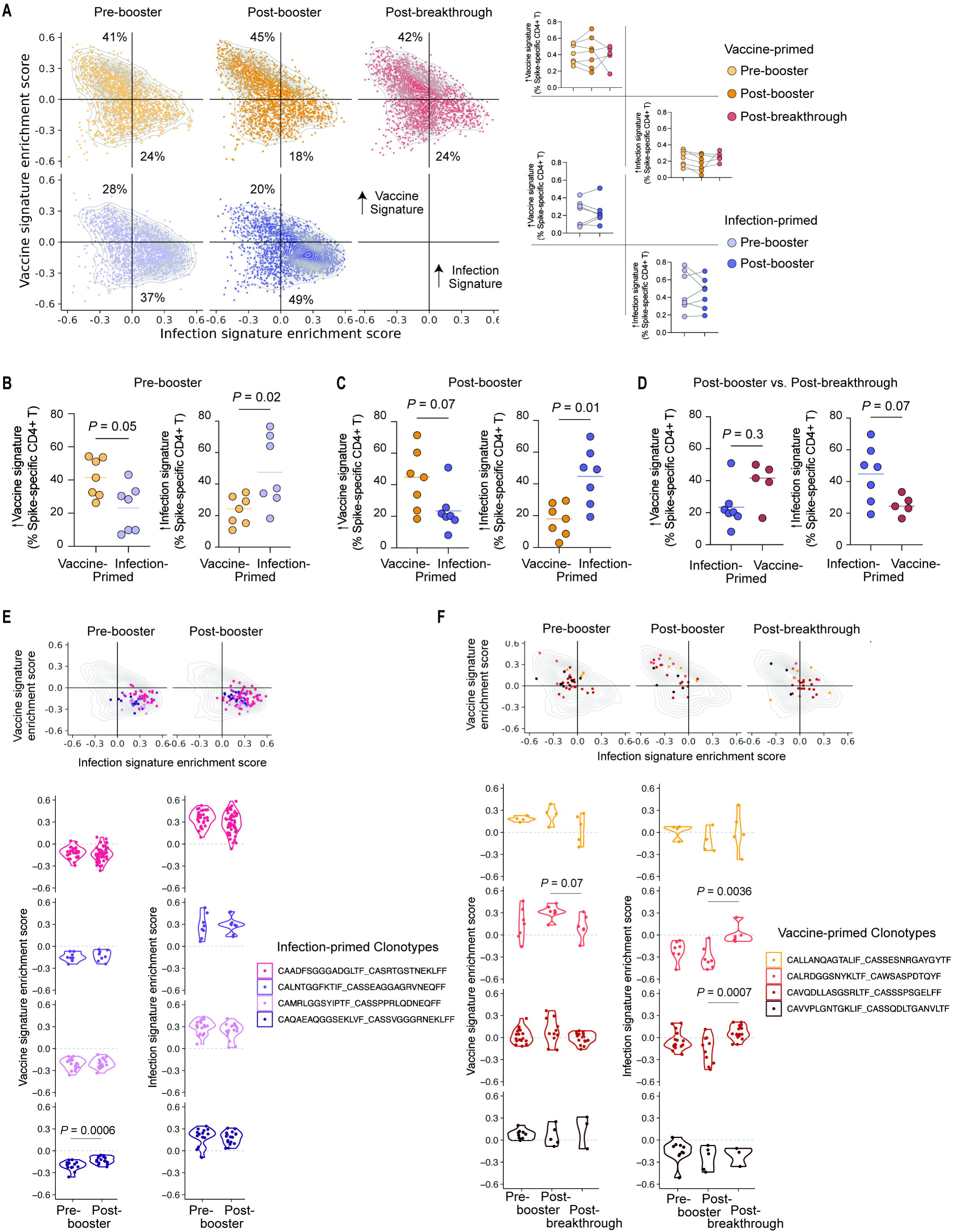
Clonotypes enriched for the vaccine-primed gene signature can undergo inflammatory imprinting by breakthrough infection. **A**. Gene set variation analysis (GSVA)-derived enrichment scores for the ‘vaccine’ and ‘infection’ signatures plotted in 2-D cartesian space for Spike-specific CD4+ T cells pooled from all individuals and split by time point and cohort. Median frequencies of first and third quadrants in black. Summary plots for each individual shown. **B-D**. Summary data at each time point. *P* values by Wilcoxon test. **E-F**. Enrichment scores for the ‘vaccine’ and ‘infection’ signatures for the four clonotypes from infection-primed participants (**E**) and vaccine-primed participants (**F**) that repeated in all time points shown. P values by Wilcoxon test. See also Figure S6.

Although breakthrough SARS-CoV-2 infection did not confer a transcriptional imprint due to inflammation in Spike-specific CD4+ T cells from vaccine-primed individuals at a bulk cell level, it remained unclear whether transcriptional profiles could change after re-exposure to Spike at a clonotypic level. To assess this, we pooled clonotypes from each cohort and identified the top four expanded clonotypes from each cohort that were present at all time points, followed by GSVA analysis for vaccine and infection signatures (**Fig 6E**). Infection-primed clonotypes were relatively homogeneous in their scores for both vaccine and infection signatures and demonstrated minimal change due to booster vaccination. Although no change was observed in the vaccine signature score for any of the four clonotypes from vaccine-primed participants following either booster immunization or breakthrough infection, two clonotypes had an increase in infection signature scores following SARS-CoV-2 breakthrough infection (**Fig. 6F**). These data suggest that re-exposure to antigen in the context of high levels of inflammation could alter the transcriptional profiles of memory CD4 T cells, albeit only weakly and in a clonotype-specific manner.

## Discussion

Memory CD4+ T cell responses are an important contributor to protective immunity, yet the molecular characteristics required for high quality memory CD4+ T cells responses are not well-understood. Using the AIM assay, we profiled the transcriptional and epigenetic landscape of Spike-specific CD4+ T cells in participants whose first exposure to Spike protein was either mRNA vaccination or SARS-CoV-2 infection. Several observations were notable. First, we uncovered a durable transcriptional signature of inflammation, which included ISGs and genes such as *GZMB*, *CCL3, CCL4, and CCL5*, in infection-primed Spike-specific CD4+ T cells relative to vaccine-primed Spike-specific CD4+ T cells. In contrast, vaccine-primed Spike-specific CD4+ T cells had enrichment for TNF/NFkB signaling pathway, Mitotic Spindle, and other pathways at both pre- and post-booster time points. Second, Spike-specific memory CD4+ T cells established by infection had distinct epigenetic landscapes likely shaped by IRF-family transcription factors. Third, inflammation due to breakthrough SARS-CoV-2 infection in vaccine-primed individuals did not grossly alter the transcriptional profile of Spike-specific memory CD4+ T cells, indicating resistance of memory CD4+ T cells to further imprinting. These observations suggest that the inflammatory context at the time of CD4+ T cell priming has lasting effects on CD4+ T cell memory.

Adaptive immune responses are the basis for long-term effective protection against pathogens, yet little is understood about why protection is durable in some scenarios and short-lived in others. Although CD4+ T cells help coordinate the immune response, the qualitative differences between different memory CD4+ T cells are not well-understood, leading to ambiguity about the optimal way to induce CD4+ T cell memory. In our studies, inflammation due to viral infection left a transcriptional signature in infection-primed Spike-specific CD4+ T cells. IFN signaling plays a central role in the immune response to COVID-19^66–69^, and indeed type I and II IFNs are likely drivers of Spike-specific cytotoxic CD4+ T cells in COVID-19 patients^51^. IFN signaling is important to T cell differentiation and proliferation^70, 71^, and consistent with this, IFNAR blockade significantly reduced granzyme-B expression in virus-specific CD4+ T cells^72^. Yet the effects of type I IFN timing and dose on memory CD4+ T cell formation and maintenance are largely unknown. Our results suggest that high levels of IFN signaling at the time of priming results in lasting effects on memory CD4+ T cell biology that may include impaired proliferative capacity, altered metabolism, and differential cytotoxicity. Future studies examining CD4+ T cell memory differentiation in the contexts of IFN signaling dose and timing are needed to provide a mechanistic understanding of the consequences of the inflammation-induced transcriptional imprinting observed here.

In our data, we observed heterogeneity in the transcriptional profiles of Spike-specific memory CD4+ T cell pools at the single-cell level in each cohort. Vaccine- and infection-derived Spike-specific CD4+ T cells demonstrated a range of functional phenotypes and exhibited heterogeneity in the molecular signatures that delineated each priming condition. This heterogeneity could be due to several factors during priming such as the kinetics of inflammation, duration (and/or order) of exposure to antigen and inflammation, and variability in level and type of inflammation in different tissue sites. Indeed, infection-primed transcriptional imprints were maintained despite repeated immunization (**Fig. 6**), whereas some, but not all, vaccine-primed clonotypes enriched for infection signature-associated transcripts after breakthrough infection. Strategies such as mitochondrial allele analysis may be needed to determine whether the enrichment was due to re-imprinting or due to *de novo* naive CD4+ T cell differentiation into memory in the context of breakthrough infection. Unraveling the intricacies of when and where memory is formed during infection or vaccination would address whether T cell heterogeneity arises throughout the course of initial exposure, over time through subsequent challenges, or with turnover of cells after repeated exposures.

To assess malleability of the transcriptional imprint, we considered re-exposure to Spike protein in high and low inflammation settings on the transcriptional profiles of vaccine- and infection-primed Spike-specific memory CD4+ T cells. Our results demonstrated that infection and vaccination induced transcriptionally distinct memory CD4+ T cell states that were sustained following subsequent exposure to Spike protein, irrespective of whether these occurred in the contexts of booster immunization or breakthrough infection. In our studies, booster immunization did not alter Spike-specific CD4+ T cell transcriptional profiles in either cohort. Moreover, vaccine-derived Spike-specific memory CD4+ T cells also largely retained their transcriptional state following breakthrough SARS-CoV-2 infection, although certain clonotypes may have been preferentially affected. These data suggest that initial priming by vaccination generated memory T cells that were relatively resistant to re-imprinting in the context of a subsequent exposure to Spike protein. Our data further suggest an epigenetic basis for the transcriptional imprinting. Indeed, six months after SARS-CoV-2 infection, Spike-specific memory CD4+ T cells had more chromatin accessibility near ISG loci, along with enrichment for IRF-family TF binding sites motifs, compared to vaccine-primed Spike-specific CD4+ T cells. Given the durability of epigenetic changes in quiescent memory T cells^73^ and the resistance to re-imprinting seen in our studies here, our data suggest persistent residual effects of the inflammatory context during CD4+ T cell priming that will have major implications for rational vaccine design.

Together, these data suggest that the imprint of inflammation during Spike-specific memory CD4+ T cell formation resulted in persistent transcriptional and epigenetic alterations which were sustained despite mRNA vaccination, relative to memory CD4+ T cells primed during vaccination. Similarly, SARS-CoV-2 breakthrough infection did not dramatically alter the transcriptional profile of vaccine-primed memory CD4+ T cells. Our results underscore the importance of prior infection history to understand CD4+ T cell immunity. Future vaccine strategies will need to consider the inflammatory state, whether directly due to the immune stimulus or indirectly due to chronic infection, at the time of CD4+ T cell memory formation.

### Limitations of the study

We found robust transcriptional differences between infection- and vaccine-primed memory CD4+ T cells, but several caveats should be considered. First, we focused on CD4+ T cells due to their role in long-term immunity^3^, but we did not examine the effects of inflammation on priming of CD8+ T cells or B cells, which also play important roles in immunity to SARS-CoV-2. Indeed, persistent ISG transcripts in Spike-specific memory CD8+ T cells were observed following recovery from COVID-19^1^. Second, we assessed circulating, but not tissue-based, memory CD4+ T cells. Tissue-resident memory CD4+ T cells play important roles in protective immunity^74^. Exploring memory CD4+ T cells in human tissue, particularly secondary lymphoid organs, would clarify whether similar transcriptional changes accrue in non-circulating cells as well. Third, better understanding of the relevance of the transcriptional differences observed here for immunity is needed, given the multifaceted nature of the immune response to re-exposure to SARS-CoV-2. Future studies including in animal models could help clarify whether the inflammatory imprinting observed here would manifest in differences in susceptibility or outcomes with re-infection. Lastly, further studies will be needed to understand the extent to which our findings generalize across age, sex, racial, and ethnic groups. These approaches will together improve our understanding of memory CD4+ T cell biology and highlight strategies for more protective vaccination.

## Supporting information

Key Resources Table

## Acknowledgements

We thank all members of NYU Vaccine Center processing and clinical staff, including Michael Tuen, Jimmy Wilson, Abdonnie Holder, Shelby Goins, Meron Tasissa, Sara Wesley Hyman, and Farzana Antara. We would like to thank the NYU Genome Technology Core for their services, with help from Paul Zappile and Emily Guzman in particular. Finally, we would like to thank all the participants who have contributed to our studies.

## Author contributions

Conceptualization: S.L.G-G and R.S.H.

Data curation: S.L.G-G

Formal analysis: S.L.G-G.

Funding acquisition: M.O., M.J.M., R.S.H.

Investigation: S. L.G.G, S.S., H.C., C.M., G.C.

Methodology: S.L.G-G., A.R.C., E.N.I., S.B.K., and R.S.H.

Project administration: R.S.H.

Supervision: R.S.H.

Validation: S.L.G-G. and R.S.H.

Visualization: S.L.G-G. and R.S.H.

Writing – original draft: S.L.G-G. and R.S.H.

Writing – review & editing: S.L.G.G, S.S., H.C., C.M., T.W., G.C., A.R.C., M.I.S., E.G., M.O., E.N.I., S.B.K., M.J.M., and R.S.H.

## STAR Methods text

### RESOURCE AVAILABILITY

#### Lead contact

Requests for further information and resources should be directed to and will be fulfilled by Ramin Herati.

#### Materials availability

This study did not generate new unique reagents.

#### Data and code availability

- Sequencing data will be deposited in GEO and dbGaP.
- All original code has been deposited on GitHub, is publicly available, and linked in the key resources table.
- Any additional information required to reanalyze the data reported in this paper is available from the lead contact upon request.

### EXPERIMENTAL MODEL AND STUDY PARTICIPANT DETAILS

#### Study design

We examined T cell responses in adults receiving a third dose of BNT162b2 vaccine at the time points indicated in Fig.1A. Following written informed consent, peripheral blood was drawn by standard phlebotomy longitudinally from 24 adults (12 vaccine-primed and 12 infection-primed) in observational studies in accordance with NYU Institutional Review Board protocols (s18-02035 and s18-02037) and previously described in^12, 41^. Participant demographics are summarized in **Tables S1-3**. Dates depicted in **Fig. S1A** and **Fig. S4A** have been randomly offset to protect patient privacy^75^.

### METHOD DETAILS

#### Blood samples processing and storage

Blood draws occurred around six months after either vaccination (“post-vaccination”) or infection (“post-infection”), eight months after second vaccination (“pre-booster”), one month post third vaccination (“post-booster”) and one month after breakthrough infection (“post-breakthrough), as depicted in Fig. 1A, Fig. 4A and Fig. 5A. Peripheral blood mononuclear cells (PBMC) were isolated from CPT vacutainers (BD Biosciences) within four hours of the blood draw and cryopreserved.

#### Activation-induced marker (AIM) analysis

Cryopreserved PBMCs were thawed and rested overnight at 37°C in RPMI 1640 with L-glutamine (Fisher) containing 10% FBS (Fisher), 2 mM L-glutamine (Fisher) and supplemented with 10 units DNaseI and 5mM MgCl_2_. The next day, cells were stimulated with combined 15-mer peptide pools encompassing the SARS-CoV-2 Spike protein (S1, S, and S+ PepTivators, Miltenyi). Sterile water was used for the unstimulated controls. After stimulation for 20 hours at 37°C, cells were washed with PBS containing 10 mM EDTA for 5 minutes prior to use in flow cytometry or exCITE-seq.

#### Flow Cytometry

Staining for flow cytometry analysis was performed using fluorescently-labeled anti-human antibodies (see Reagent Table, **STAR Methods**). Briefly, cells following AIM assay underwent Fc blockade with Human TruStain FcX (BioLegend) and NovaBlock (Thermo Fisher Scientific) for 10 min at room temperature, followed by surface antibody staining at room temperature for 20 min in the dark. Samples not planned for use in scRNAseq were permeabilized using the eBioscience Foxp3/Transcription Factor Staining Buffer Set (ThermoFisher) for 20 minutes at room temperature in the dark. Following permeabilization, cells were intracellularly stained for 1 hour at room temperature in the dark, followed by resuspension in 1% para-formaldehyde (Electron Microscopy Sciences). All samples were acquired on a five-laser Aurora cytometer (Cytek Biosciences).

#### Tetramer staining

PBMCs used for tetramer staining were thawed and rested overnight at 37°C in RPMI 1640 with L-glutamine (Fisher) containing 10% FBS (Fisher), 2 mM L-glutamine (Fisher) and supplemented with DNase and MgCl_2_. The next day, cells were stained with Fc blockade at 1:100 and BV421-labeled HLA-DPB1*04:01 S_167-180_ tetramer (MBL) at 1:20 for 2 hours at 37°C, followed by surface antibody staining, permeabilization, intracellular staining, and acquisition as described above.

#### Expanded cellular indexing of transcriptomes and epitopes by sequencing

Following overnight stimulation in the AIM assay (above), cells were stained with antibodies against CD69 and CD137 conjugated to PE-Dazzle594 and PE, respectively (Biolegend), for 30 minutes at room temperature in the dark, followed by magnetic bead enrichment using the EasySep Human PE Positive Selection Kit (STEMCELL Technologies). Cells were processed for expanded cellular indexing of transcriptomes and epitopes by sequencing (exCITE-seq) using Chromium Next GEM Single Cell 5’ HT Kit v2 (10X Genomics). Cells were stained with hashtag oligos (Biolegend) as previously described^45, 76–78^. Cells were pooled and loaded onto Chromium HT Chips and run on a Chromium controller (10X Genomics). Gene expression, V(D)J, and surface protein expression libraries were made using the 5’ Feature Barcode Kit, Chromium Single Cell V(D)J Amplification Kit, and Chromium Next GEM Single Cell 5’ Library Kit (10X Genomics) as recommended by the manufacturer. Libraries were pooled and sequenced using the NovaSeq 6000.

#### exCITE-seq data processing

Cellranger software v7 (10X Genomics) was used to align FASTQ files to the human genome (GRCh38 ensemble) and to perform protein expression evaluation. Seurat v4.3.0^46^ was used to process single cell libraries and integrate all the exCITE-seq modalities. Empty droplets were excluded based on the barcode-rank distribution. Following CLR normalization, HTOs were demultiplexed using the Seurat function *HTODemux()*. Cells were excluded based on mitochondrial content >8% or if the cell contained fewer than 400, or more than 7000, UMI counts. Doublets were identified using scDblFinder^79^ based on an estimated doublet rate of 0.4%. RNA was normalized across batches using SCTransform and filtered counts were integrated in Seurat using *SelectIntegrationFeatures()* for 3000 features, followed by *FindIntegrationAnchors()* using the first 30 dimensions with ‘rpca’ dimensionality reduction and k.anchors=20. TCR sequences were processed and analyzed using scRepertoire v1.7.4^80^. SPICE was used to analyze polyfunctionality (version 6.1)^81^. The following list of genes^82–84^ was used to identify functional phenotypes in the AIM-Reactive CD4+ T cluster: *IFNG*, *TNF*, *IL2*, *IL12A*, *CXCR3*, *CCR5*, *STAT4*, *TBX21*, *RUNX3*, *IL4*, *IL5*, *IL13*, *CXCR4*, *GATA3*, *STAT6*, *CCR4*, *IL21*, *IL17A*, *RORC*, *STAT3*, *CCR6*, *IL10*, *IL2RA*, *FOXP3*, *CCR7*, *TGFB1*, *CXCR5*. GO was performed using Metascape^85^. Differential expression analysis was performed using DESeq2 v1.34.0^86^ after filtering for genes with at least 10 counts in at least 3 samples. Pre-ranked gene set enrichment analysis (GSEA) was performed with 10,000 permutations of gene sets from the MSigDB.

#### Transcriptomic, epitope, and accessibility by sequencing

Following overnight stimulation in the AIM assay (above), CD4+ T cells were isolated using the EasySep Human CD4+ T Cell Isolation Kit (STEMCELL Technologies). Next, isolated CD4+ T cells were stained with antibodies against CD69 and CD137 conjugated to PE-Dazzle594 and PE, respectively (Biolegend), and with hashtag oligos (Biolegend), for 30 minutes at room temperature in the dark, followed by magnetic bead enrichment using the EasySep Human PE Positive Selection Kit (STEMCELL Technologies). Cells were then processed for transcriptomic, epitope, and accessibility measurement (TEA-seq)^55^ using the Chromium Next GEM Single Cell Multiome ATAC + Gene Expression Kit (10XGenomics). Cells were pooled and loaded onto Chromium J Chips and run on a Chromium X controller (10XGenomics). Gene expression and ATAC libraries were made using the Chromium Next GEM Single Cell Multiome ATAC Kit A, Chromium Next GEM Single Cell Multiome Reagent Kit A, Library Construction Kit (10XGenomics) following the protocols recommended by the manufacturer. Surface protein expression libraries were made as detailed in the TEA-seq protocol. Libraries were pooled at desired concentrations and sequenced using the NovaSeq 6000.

#### TEA-seq data processing

RNA and ATAC libraries were aligned using cellranger-arc v1.2.0 (10XGenomics) against the human genome (GRCh38 ensemble). Primary data analysis and statistical analysis were then performed using the R environment. BarCounter^55^, Seurat v4.3.0^46^ and Signac v1.9.0^87^ were used to process single cell libraries and integrate all the TEA-seq modalities. HTOs were demultiplexed using *HTODemux()*, and scDblFinder identified doublets^79^ based on an estimated doublet rate of 0.8%. RNA was normalized across batches using the same integration methods applied to exCITE-seq data. Pre-ranked GSEA was performed with 10,000 permutations of gene sets from the MSigDB. To create a merged peak list, OCRs originally called by cellranger-arc were normalized across batches in Seurat using *IntegrateEmbeddings()*. Subsetting down to the AIM-Reactive CD4+ T cluster, peaks were recalled using MACS2^88^. Differential accessibility was performed using the Seurat *FindMarkers()* function with test.use = ‘LR’. We identified peaks which contained DNA motifs that were present in the JASPAR database for humans^89^. The Signac library’s *FindMotifs()* function was used to identify overrepresented motifs in the top DARs in each cohort (nominal *P* < 5×10^-3^) compared to background, matched for overall GC content. ChEA3^64^ was used to predict TF activity associated with input gene lists (**Table S4**).

### QUANTIFICATION AND STATISTICAL ANALYSIS

#### Statistics

Nonparametric tests were preferentially used throughout using two-tailed tests at α=0.05, unless otherwise indicated. Where paired analyses were performed, the unpaired data points were excluded. Outlier analysis was not performed and thus outliers were not excluded from the primary analyses. Genes were considered differentially expressed at a false discovery rate (FDR) <= 0.05. Prism 9.0 and R were used to perform statistical analyses. Study schematics were made using BioRender.

