## Supplementary material for "Inflammation durably imprints memory CD4+ T cells": Key Resources Table

| REAGENT or RESOURCE | SOURCE | IDENTIFIER |
|---|---|---|
| **Antibodies** | | |
| Anti-human TNF-a (MAb11) | BD Biosciences | Cat# 563996 |
| Anti-human CD19 (SJ25C1) | Biolegend | Cat# 612939 |
| Anti-human CD25 (2A3) | BD Biosciences | Cat# 612919 |
| Anti-human CD14 (M5E2) | BD Biosciences | Cat# 612902 |
| Anti-human CD4 (SK3) | Biolegend | Cat# 344674 |
| Anti-human CD56 (5.1H11) | Biolegend | Cat# 362539 |
| Anti-human CCR7 (G043H7) | Biolegend | Cat# 353224 |
| Anti-human CD27 (O323) | Biolegend | Cat# 67-0279-42 |
| Anti-human CD137 (4B4-1) | BD Biosciences | Cat# 747353 |
| Anti-human CD71 (OKT9) | Invitrogen | Cat# 78-0719-42 |
| Anti-human OX40 (ACT-35) | Invitrogen | Cat# 11-1347-41 |
| Anti-human CD3 (SK7) | Biolegend | Cat# 344851 |
| Anti-human PD-1 (EH12.1) | BD Biosciences | Cat# 566460 |
| Anti-human GZMB (GB11) | BD Biosciences | Cat# 568705 |
| Anti-human CD69 (FN50) | Biolegend | Cat# 310942 |
| Anti-human CD8 (SK1) | Biolegend | Cat# 344761 |
| Anti-human CD45RA (MEM-56) | Fisher Scientific | Cat# MHCD45RA18 |
| Anti-human CD200 (OX-104 ) | Biolegend | Cat# 329211 |
| Anti-human CXCR3 (G025H7) | Biolegend | Cat# 353760 |
| Anti-human CCR6 (G034E3) | Biolegend | Cat# 353404 |
| Anti-human IFNg (4S.B3) | Biolegend | Cat# 502551 |
| Anti-human ICOS (C398.4A) | Biolegend | Cat# 313536 |
| Anti-human CD38 (HIT2) | Biolegend | Cat# 303549 |

| | | |
|---|---|---|
| TotalSeq™-C0252 anti-human Hashtag 2 Antibody | Biolegend | Cat# 394663 |
| TotalSeq™-C0253 anti-human Hashtag 3 Antibody | Biolegend | Cat# 394665 |
| TotalSeq™-C0254 anti-human Hashtag 4 Antibody | Biolegend | Cat# 394667 |
| TotalSeq™-C0255 anti-human Hashtag 5 Antibody | Biolegend | Cat# 394669 |
| TotalSeq™-C0256 anti-human Hashtag 6 Antibody | Biolegend | Cat# 394671 |
| TotalSeq™-C0257 anti-human Hashtag 7 Antibody | Biolegend | Cat# 394673 |
| TotalSeq™-C0258 anti-human Hashtag 8 Antibody | Biolegend | Cat# 394675 |
| TotalSeq™-C0259 anti-human Hashtag 9 Antibody | Biolegend | Cat# 394677 |
| TotalSeq™-A0251 anti-human Hashtag 1 Antibody | Biolegend | Cat# 394601 |
| TotalSeq™-A0252 anti-human Hashtag 2 Antibody | Biolegend | Cat# 394603 |
| TotalSeq™-A0253 anti-human Hashtag 3 Antibody | Biolegend | Cat# 394605 |
| TotalSeq™-A0254 anti-human Hashtag 4 Antibody | Biolegend | Cat# 394607 |
| TotalSeq™-A0256 anti-human Hashtag 6 Antibody | Biolegend | Cat# 394611 |
| TotalSeq™-A0257 anti-human Hashtag 7 Antibody | Biolegend | Cat# 394613 |
| TotalSeq™-A0258 anti-human Hashtag 8 Antibody | Biolegend | Cat# 394615 |
| TotalSeq™-A0259 anti-human Hashtag 9 Antibody | Biolegend | Cat# 394617 |
| **Biological samples** | | |
| Adult Human Peripheral Blood Samples | NYU Langone Vaccine Center | |
| **Chemicals, peptides, and recombinant proteins** | | |
| Lymphoprep | STEMCELL | Cat# 07851 |
| PepTivator SARS-CoV-2 Prot_S | Miltenyi Biotec | Cat# 130-126-701 |
| PepTivator® SARS-CoV-2 Prot_S1 | Miltenyi Biotec | Cat# 130-127-048 |
| PepTivator SARS-CoV-2 Prot_S+ | Miltenyi Biotec | Cat# 130-127-312 |
| BV421-labeled HLA-DPB1*04:01 $S_{167-180}$ (TFEYVSQPFLMDLE) | MBL International | Cat# TBCM4-P1041I-4 |
| LIVE/DEAD Fixable Dead Cell Stain Kit | Invitrogen | Cat# L23105 |
| Anti-human TruStain Fc-X | BioLegend | Cat# 422301 |

| | | |
|---|---|---|
| NovaBlock | ThermoFisher | Cat# B001T03F01 |
| Brilliant Staining Buffer | BD Biosciences | Cat# 563794 |
| FcR Blocking Reagent, human | Miltenyi Biotec | Cat# 130-059-901 |
| PFA | Electron Microscopy Sciences | Cat# 15714-S |
| RPMI 1640 Medium 1X with L-Glutamine | ThermoFisher | Cat# 10040CM |
| Monensin | BD Biosciences | Cat# 554724 |
| RNase Inhibitor | Sigma-Aldrich | Cat# 3335399001 |
| Digitonin | Sigma-Aldrich | Cat# D141-100MG |
| **Critical commercial assays** | | |
| EasySep Human CD4+ T Cell Isolation Kit | STEMCELL | Cat# 17952 |
| EasySep Release Human PE Positive Selection Kit | STEMCELL | Cat# 17654 |
| NovaSeq 6000 S4 Reagent Kit v1.5 (200 Cycles) | Illumina | Cat# 20028313 |
| eBioscience Intracellular Fixation & Permeabilization Buffer Set | ThermoFisher | Cat# 88-8824-00 |
| **Deposited data** | | |
| exCITEseq for longitudinal sampling of infection- and vaccine-primed participant PBMCs after AIM assay | This paper | Data will be deposited in GEO |
| TEAseq for infection- and vaccine-primed participant PBMCs after AIM assay | This paper | Data will be deposited in dbGaP |
| High Confidence SARS-CoV-2 TCRB Sequences | Adaptive Biotechnologies | https://clients.adaptivebiotech.com/pub/covid-2020 |
| **Software and algorithms** | | |
| FlowJo (Version: 10.8.1) | BD Life Sciences | https://www.flowjo.com/ |
| GraphPad Prism 9 | GraphPad | https://www.graphpad.com/ |
| SPICE 6 | Ref [81] | https://niaid.github.io/spice/ |
| GSEA 4.2.3 | Ref [54] | https://www.gsea-msigdb.org/ |
| Metascape | Ref [85] | https://metascape.org/ |

| | | |
|---|---|---|
| ChEA3 | Ref [64] | https://maayanlab.cloud/chea3/ |
| R Version 4.1.2 | R Foundation for Statistical Computing | https://www.r-project.org/ |
| Bcl2fastq | Illumina | https://support.illumina.com |
| BarCounter | Ref [55] | https://github.com/AllenInstitute/aifi-swanson-teaseq |
| Seurat v4.3.0 | Ref [46] | |
| Signac v1.9.0 | Ref [87] | |
| Cell Ranger Multi Version 7 | 10XGenomics | https://www.10xgenomics.com/ |
| Cell Ranger ARC Version v1.2.0 | 10XGenomics | https://www.10xgenomics.com/ |
| Custom bioinformatics scripts | N/A | https://github.com/teamTfh/CD4T_DifferentialPriming |
| **Other** | | |
| Graphical abstract and workflow schematics | This paper | BioRender |
