## Supplemental Figures and Tables S1-3 for "Inflammation durably imprints memory CD4+ T cells"

**Affiliations**:
[1]Department of Medicine, New York University Grossman School of Medicine; New York, NY, USA.
[2]Department of Microbiology, New York University School of Medicine; New York, NY, USA.
[3]Department of Pathology, New York University School of Medicine; New York, NY, USA.

***Corresponding author and lead contact**:
Ramin Sedaghat Herati


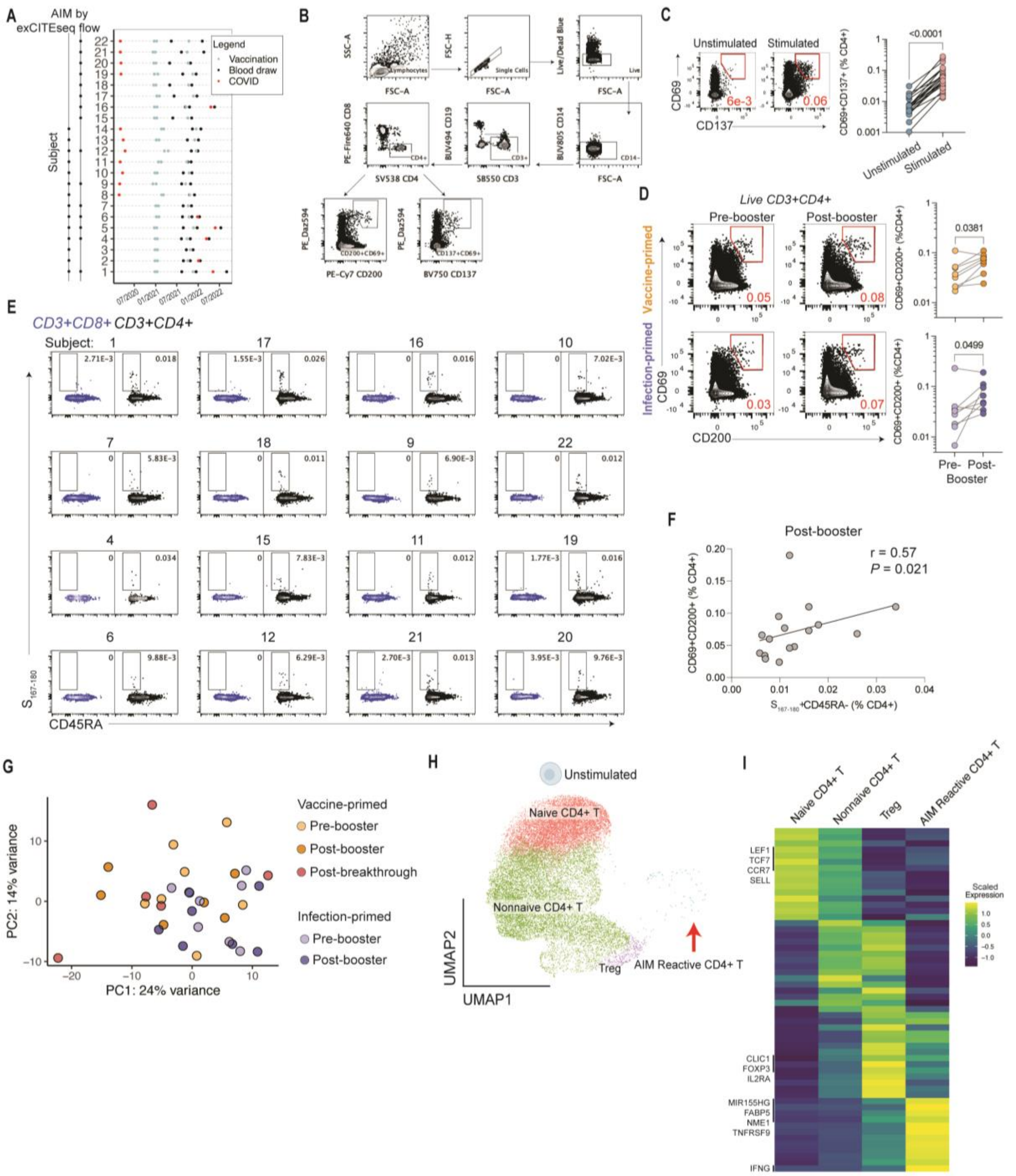


**Figure S1. Related to Figure 1**

**A**. Sample timeline for each participant. Black dots indicate PBMC samplings which were used in this study. **B**. Gating scheme. **C**. Example flow plots of unstimulated and stimulated samples showing expression of CD69+ CD137+ co-expression in CD4+ cells ($P < 10^{-4}$, Wilcoxon matched-pairs signed rank test). **D**. Example flow plots for vaccine- and infection-primed participants at pre-booster and post-booster time points for expression of CD69 and CD200 after AIM assay. Frequency shown in red. Summary data for vaccine-primed (orange, n = 9) and infection-primed (purple, n=9) participants. *P* values by Wilcoxon matched-pairs signed rank test. **E**. $S_{167-180}$+ tetramer staining in CD8+ (blue, negative control) and CD4+ (black) cells for all participants. **F**. Correlation between frequency of CD69+ CD200+ CD4+ T cells after AIM and the frequency of $S_{167-180}$+ tetramer at the post-booster time point (n = 16). **G**. PCA of all post-AIM PBMC libraries. **H**. UMAP of all unstimulated CD4+ T cells pooled across samples and clustered for gene expression. **I**. Scaled expression of genes differentially expressed in each cluster at adjusted $P < 0.05$.

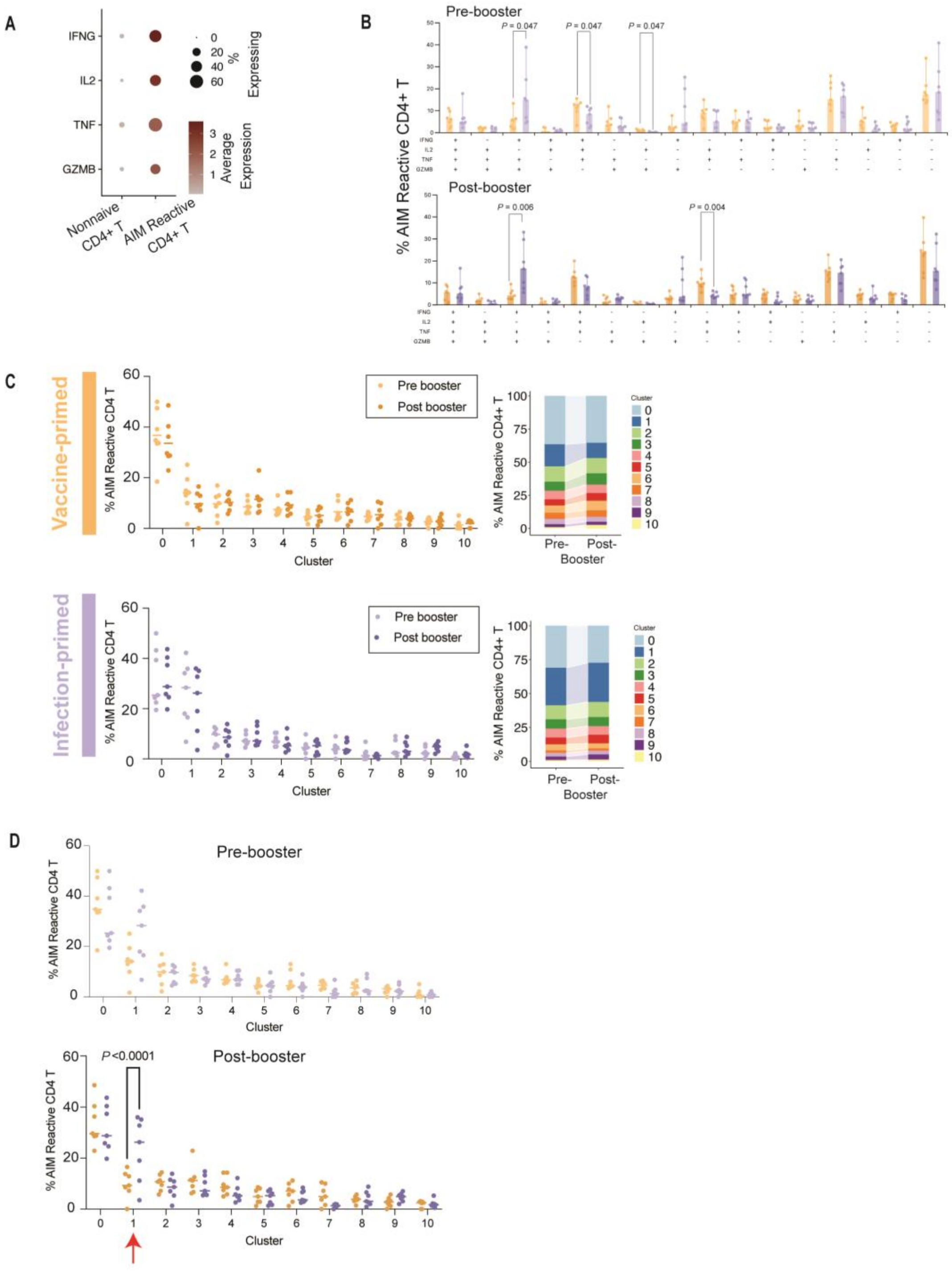


**Figure S2. Related to Figure 2**

**A**. Proportion of cells expressing transcripts for *IFNG*, *IL2*, *TNF*, and *GZMB*, and mean expression of *IFNG*, *IL2, TNF*, and *GZMB*. **B**. Polyfunctionality frequencies plotted for each sample. *P* values by Wilcoxon test. **C**. Median cluster distribution at pre- and post-booster time points for vaccine-primed and infection-primed cohorts. **D**. Individual participant cluster distribution at pre- and post- booster timepoints comparing vaccine- versus infection- primed Spike-specific CD4+ T cells. *P* values by paired two-way ANOVA with Sidak posttest.

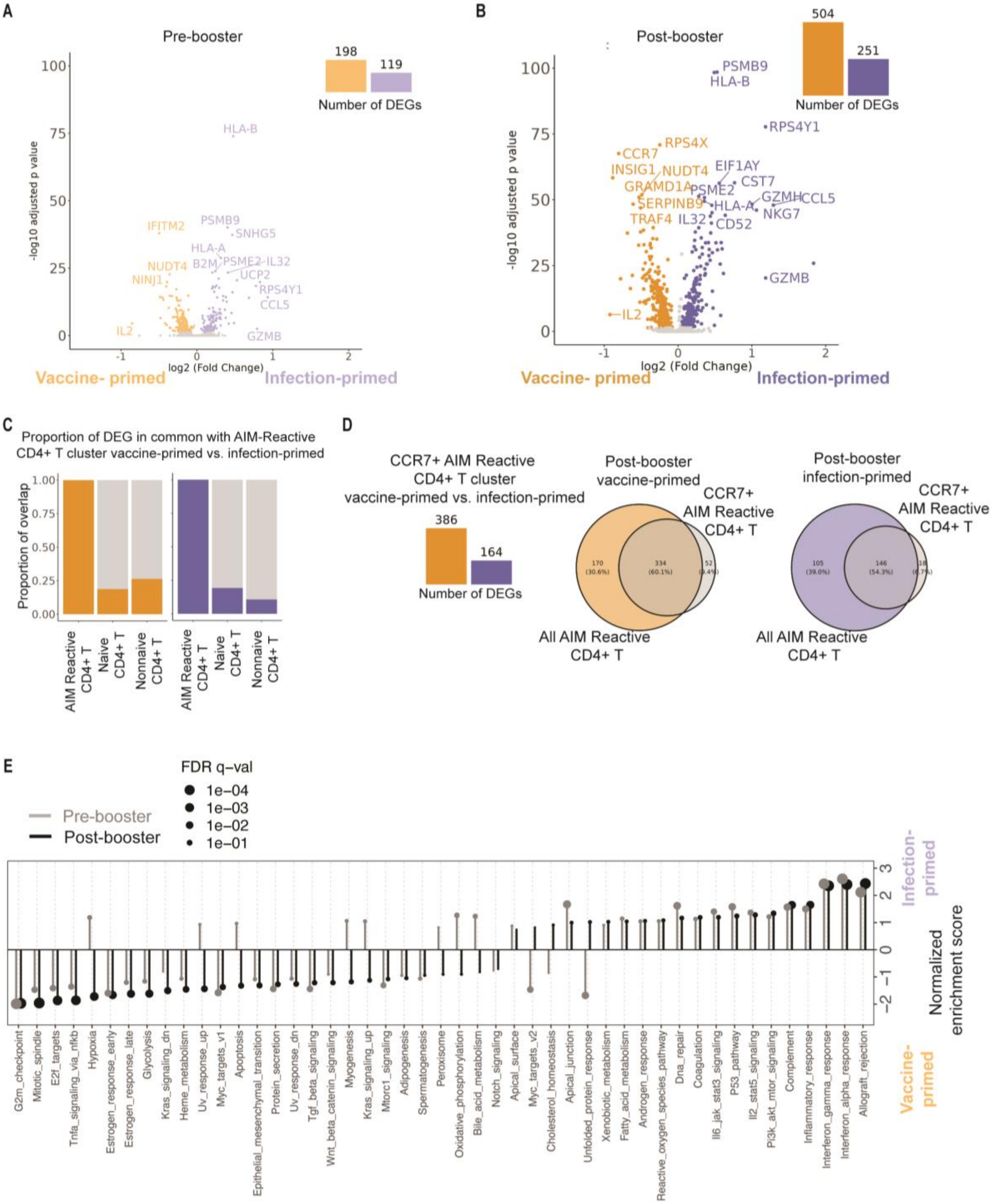


**Figure S3. Related to Figure 3 and Tables S4-S5**

**A-B.** Volcano plots of differentially expressed genes at pre-booster (**A**) and post-booster (**B**) time points. Adjusted *P* values reported. Genes in purple denote enrichment in infection-primed Spike-specific CD4+ T cells and orange for vaccine-primed Spike-specific CD4+ T cells. **C**. DEG shared between the vaccine-primed (orange) and infection-primed (purple) DEGs from

AIM-Reactive CD4+ T cells and the indicated CD4+ T cell clusters for the post-booster time point. **D**. Overlap of DEGs in CCR7+ AIM-Reactive CD4+ T cells between vaccine- and infection-primed cells and all AIM-Reactive CD4+ T cells for the post-booster time point. **E**. GSEA results for Hallmark gene sets at both pre- (gray) and post- (black) booster time points. Positive enrichment scores denote enrichment towards the infection-primed cohort.

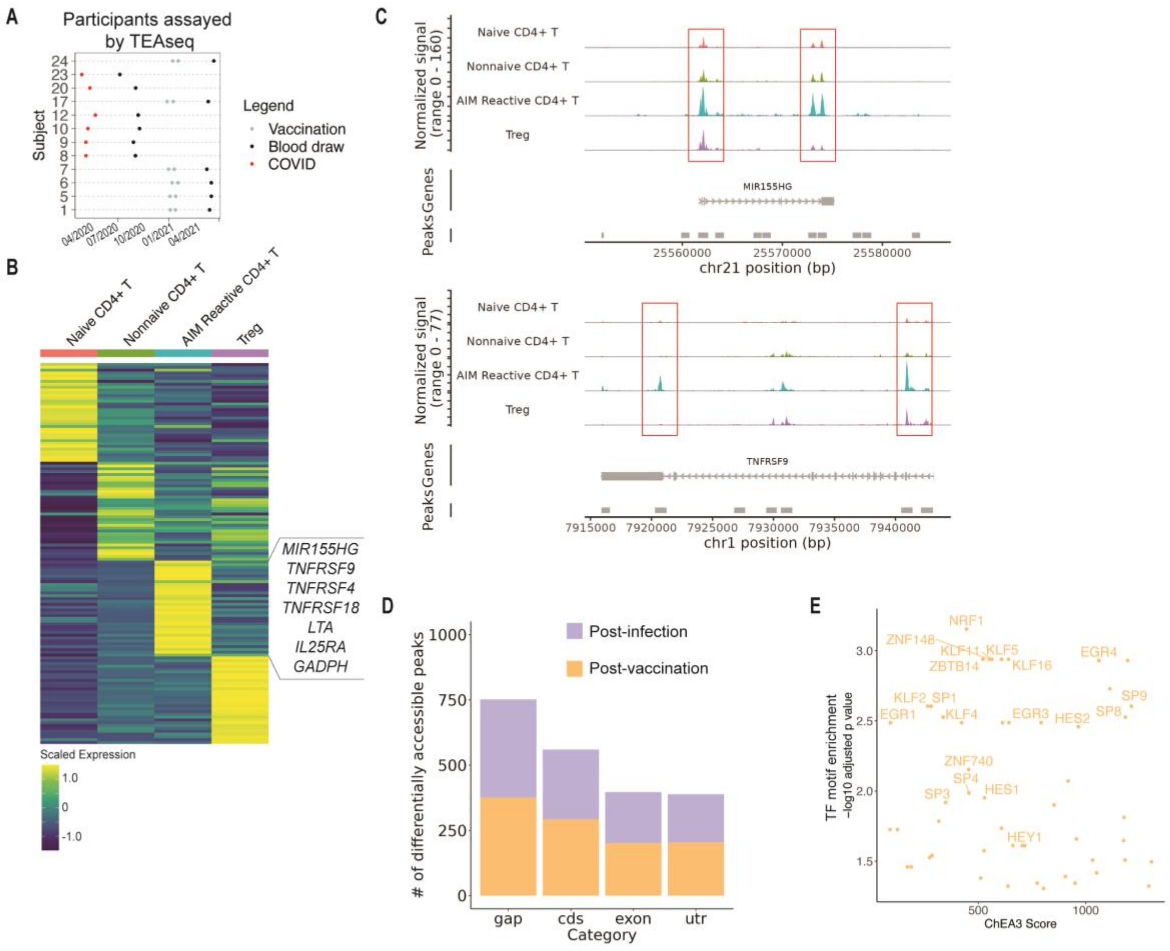


**Figure S4**. **Related to Figure 4 and Tables S6-S7**

**A**. Sample timeline for each participant. **B**. Scaled expression of genes differentially expressed in each cluster at adjusted $P < 0.05$. **C**. Representative ATAC-seq tracts shown at the *MIR155HG* and *TNFRSF9* loci in each cluster. **D**. Bar graph showing distribution of ATAC-seq peaks by cohort. **E**. TFs shown for the $-\log_{10}$ adjusted $P$ value from post-vaccination OCR motif analysis against the TF predicted by ChEA3 using the vaccine-primed gene expression data.

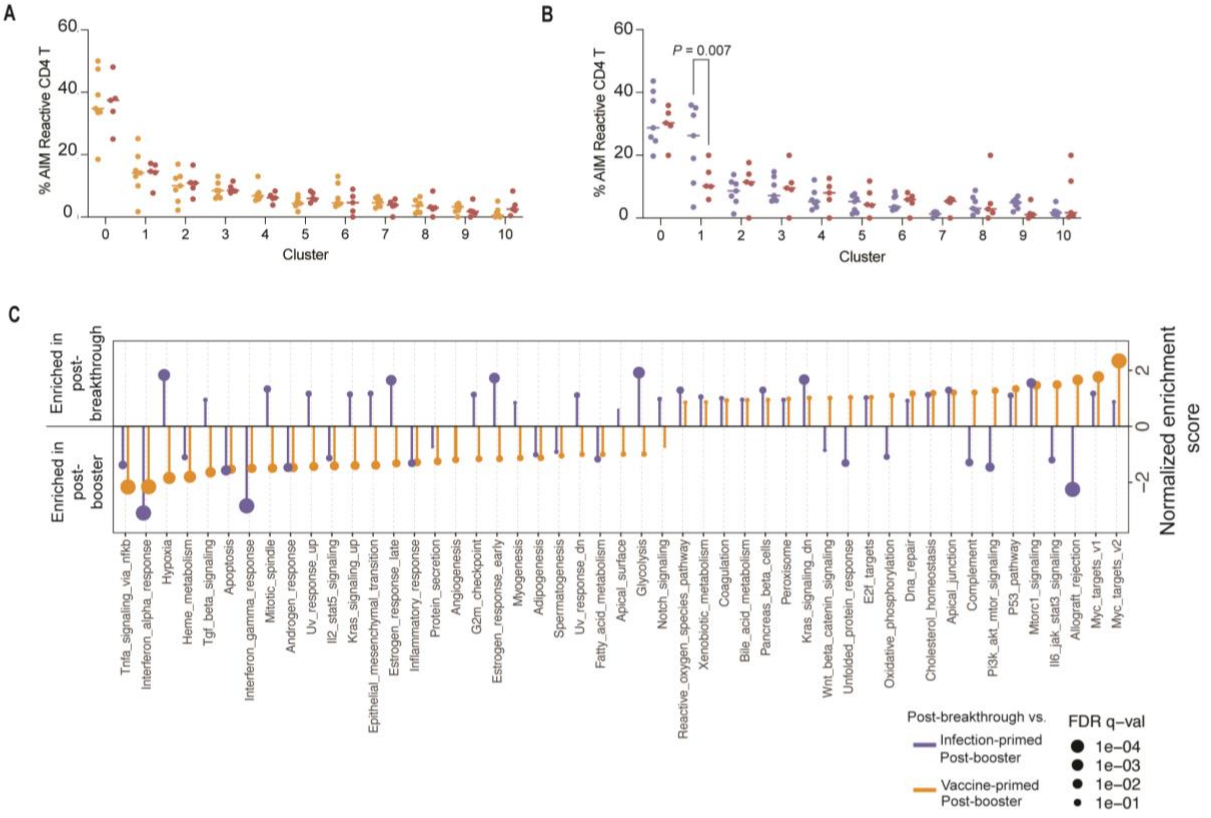


**Figure S5. Related to Figure 5 and Tables S8-S9**

**A-B.** Cluster abundance for individual samples at post-booster time points for vaccine- (**A**) and infection-primed (**B**) cohorts compared to post-breakthrough responses. *P* value by unpaired two-way ANOVA with Sidak's post-test. **C**. GSEA for Hallmark gene sets. Positive enrichment scores denote enrichment for the post-breakthrough samples.

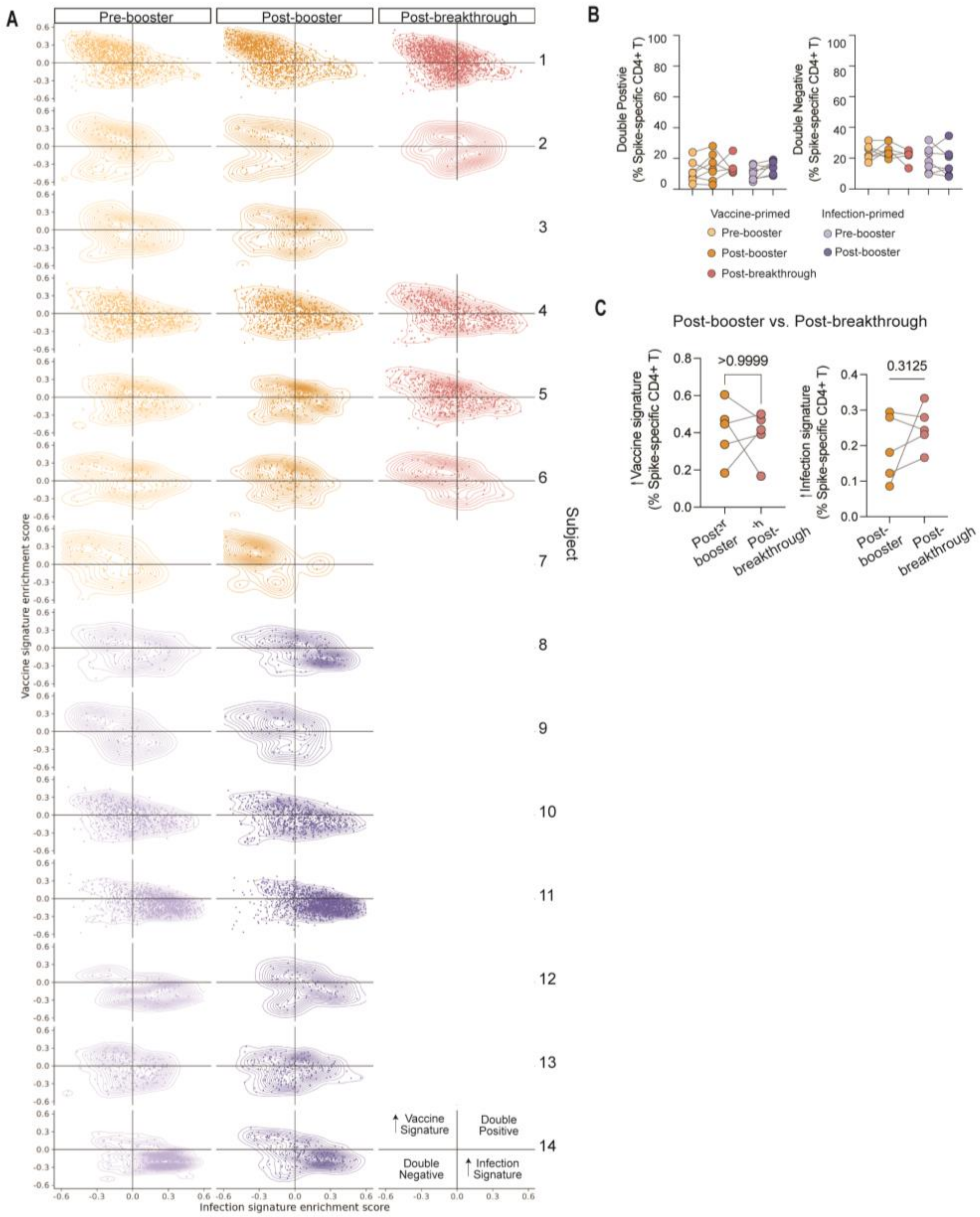


**Figure S6. Related to Figure 6**

**A**. Gene set variation analysis (GSVA)-derived enrichment scores for the 'vaccine' and 'infection' signatures shown as one row per individual and time point. **B**. Summary plots for double negative and double positive quadrants. **C**. Paired comparisons of summary data from (6A) from vaccine-primed participants post-booster and post-breakthrough. *P* values by Wilcoxon matched-pairs signed rank test.

**Table S1. Participant demographics, Related to Figure 1**

| | **Vaccine-primed** | **Infection-primed** |
|---|---|---|
| **Number of participants** | 12 | 12 |
| **Age** | | |
| Median | 39 | 42.5 |
| Range | 28 – 62 | 32 – 54 |
| **mRNA Vaccine Type** | | |
| Moderna | 0 | 3 |
| Pfizer | 12 | 9 |
| **Sex (% Female)** | 50% | 41.7% |
| **Days between onset of COVID-19 symptoms and third vaccine dose** | | |
| Median | | 588 |
| Range | | 566 - 650 |
| **Days between third vaccine dose and onset of breakthrough COVID-19 symptoms** | | |
| Median | 167 | |
| Range | 75 - 187 | |
| **Race (%)** | | |
| **White or Caucasian** | 83.3% | 75% |
| **Asian** | 16.7% | 8.3% |
| **Black or African-American** | 0% | 0% |
| **Other** | | 16.7% |
| **Hispanic (%)** | 0% | 8.3% |

**Table S2. Clinical COVID-19 disease for infection-primed participants, Related to Figure 1**

| Participant | Number of days between onset of COVID-19 symptoms and first dose of vaccine | Number of days between onset of COVID-19 symptoms and third dose of vaccine | COVID-19 diagnostic test | World Health Organization COVID-19 Severity Score | Days Hospitalized | Outcome |
|---|---|---|---|---|---|---|
| 8 | 360 | 606 | NAAT | 7 | 17 | Required intubation, treated with immunomodulator, improved and was discharged from hospital |
| 9 | 284 | 566 | NAAT | 2 | 0 | |
| 10 | 279 | 572 | NAAT | 2 | 0 | |
| 11 | 329 | 573 | NAAT | 2 | 0 | |
| 12 | 338 | 610 | Commercial antibody test | 2 | 0 | |
| 13 | 284 | 614 | Commercial antibody test | 2 | 0 | |
| 14 | 353 | 650 | Commercial antibody test | 2 | 0 | |
| 19 | 286 | 615 | None | 2 | 0 | |
| 20 | 281 | 573 | PCR | 2 | 0 | |
| 21 | 280 | 573 | PCR | 2 | 0 | |
| 22 | 290 | 588 | Rapid Antigen Test | 2 | 0 | |
| 23 | 460 | N/A | PCR | 2 | 0 | |

**Table S3. Clinical COVID-19 disease for participants with breakthrough infections, Related to Figure 5**

| Participant | Number of days between third dose of vaccine and onset of COVID-19 symptoms | COVID-19 diagnostic test | World Health Organization COVID-19 Severity Score |
|---|---|---|---|
| 1 | 172 | NAAT | 2 |
| 2 | 93 | NAAT | 2 |
| 4 | 167 | NAAT | 2 |
| 5 | 187 | Rapid Antigen Test | 2 |
| 6 | 75 | NAAT | 2 |
